# Vascular HIF2 signaling prevents cardiomegaly, alveolar congestion and capillary remodeling during chronic hypoxia

**DOI:** 10.1101/2024.09.03.610947

**Authors:** Albendea-Gomez Teresa, Mendoza-Tamajon Susana, Castro-Mecinas Rosana, Escobar Beatriz, Rocha Susana Ferreira, Urra-Balduz Sonia, Nicolas-Avila Jose Angel, Oliver Eduardo, Villalba-Orero Maria, Martin-Puig Silvia

## Abstract

Hypoxia is associated with the onset of cardiovascular diseases including cardiac hypertrophy and pulmonary arterial hypertension (PAH). Endothelial HIF2 signaling mediates pulmonary arterial remodeling and subsequent right ventricular systolic pressure (RVSP) elevation during chronic hypoxia, encouraging novel therapeutic opportunities for PAH based on specific HIF2 inhibitors. Nevertheless, HIF2 relevance beyond the pulmonary endothelium or in the cardiac adaptation to hypoxia remains elusive. Wilms tumor 1 lineage contributes to heart and lung vascular compartments including pericytes, endothelial and smooth muscle cells. Here we describe the response to chronic hypoxia of a novel HIF2 mutant mouse model in the Wt1 lineage (*Hif2/Wt1* cKO). *Hif2/Wt1* cKO is protected against pulmonary remodeling and increased RVSP induced by hypoxia, but displays alveolar congestion, inflammation and hemorrhages associated with microvascular instability. Furthermore, lack of HIF2 in the Wt1 lineage leads to cardiomegaly, capillary remodeling, right and left ventricular hypertrophy, systolic dysfunction and left ventricular dilation, suggesting pulmonary-independent cardiac direct roles of HIF2 in hypoxia. These structural defects are partially restored upon reoxygenation, while functional parameters remain altered. Our results suggest that cardiopulmonary HIF2 signaling prevents excessive vascular proliferation during chronic hypoxia and define novel protective roles of HIF2 to warrant stable microvasculature and organ function.

## Introduction

Low oxygen tensions induce the activation of HIFs that are heterodimeric transcription factors composed by an oxygen-regulated α subunit, and a constitutively expressed oxygen-independent β subunit, also known as aryl hydrocarbon receptor nuclear translocator (ARNT) (1). In normal oxygen conditions or normoxia, the proline residues of HIFα subunits are hydroxylated by oxygen-dependent prolyl-4-hydroxylases (PHDs). Von Hippel–Lindau protein (VHL), binds to the hydroxylated HIFα and acts as a substrate recognition component of the E3 ubiquitin ligase complex, which leads to proteosomal degradation of HIFα protein. Under hypoxia, the activity of PHDs is suppressed, and HIFα subunits translocate into the nucleus to bind HIF1β. Then, the heterodimer HIFα/HIFβ binds to the hypoxia response elements (HREs) in its target genes, resulting in their transcriptional upregulation (2-4). HIF1 is known to be associated with the upregulation of glycolytic genes such as glucose transporter 1 (GLUT1), phosphoglycerate kinase (PGK) or lactate dehydrogenase A (LDHA), which function to metabolically adapt the tissue to oxygen deprivation and anaerobic ATP synthesis. HIF2 induces erythropoietin (EPO) and vascular endothelial growth factor (VEGF), which are important to improve oxygen supply to the hypoxic region (5, 6). Although HIF1 and HIF2 bind to an identical core-binding motif within the HRE: 5′-RCGTG-3′, they have unique targets not compensated by the complementary isoform. HIF1 contributes more to the acute hypoxia-driven transcriptional responses, like cellular glycolysis or adenosine release after injury, while HIF2 has been mostly related to chronic adaptation to hypoxia (7). Both isoforms have been related with several pathologies.

It is well established that HIFs play an important role in the heart homeostasis and along the progression of cardiovascular diseases (8). During development, HIF1α is expressed in the embryonic heart (9-12), where it governs a glycolytic metabolism of the compact myocardium (11, 12). In contrast, the role of HIF2 during heart development remains poorly understood. Global HIF2 deletion affects catecholamine production in the organ of Zuckerkandl, leading to prominent bradycardia and cardiac dysfunction (13). Nevertheless, the phenotype of full HIF2 knock out mice varies depending on the genetic background (13-15). In the lung, HIF signaling have also been reported to execute important functions. HIF1α is implicated in bronchial epithelial formation, while HIF2 is mostly expressed in vascular endothelium and alveolar type II cells, playing a crucial role in vascular morphogenesis and surfactant production during lung development (16). In contrast to the limited knowledge about HIF2 function in the heart, the role of HIF2 signaling has been extensively studied during the development and progression of pulmonary arterial hypertension (PAH). PAH is a cardiovascular disorder that can appear in patients with chronic obstructive pulmonary disease (COPD), after prolonged exposure to hypoxia and/or because living at high altitude (World Health Organization Class 3) (16). PAH is characterized by significant vascular remodeling of the distal pulmonary arteries resulting in reduced vascular lumen (17) and increased resistance, which leads to elevation of the right ventricular systolic pressure (RVSP) that eventually could lead to heart failure (HF) (16-20). In this context, the role of HIF1 and HIF2 in endothelial cells (ECs) during the onset of PAH has been extensively studied. Several lines of evidences using *VE-Cadherin-Cre (Cdh5)* (19), *Tie2-Cre* (20), or *L1Cre* (18) models to mediate endothelial-specific deletion of HIFs have uncovered the essential role of HIF2 in mediating pulmonary arterial remodeling associated to vasoconstriction, proliferation of vascular smooth muscle cells (VSMC) and ECs and fibrosis, causing the occlusion of the pulmonary arteries and increasing the RVSP(21). Recent publications show the relevance of HIF signaling during muscularization in mural cells, either VSMC (17, 22) or pericytes (PCs) (23). However, the role of HIF2 in other cell types within the vascular compartment remains elusive.

Wilms tumor 1 (Wt1) is a transcription factor with a critical role in organogenesis and adult homoeostasis. During embryogenesis Wt1 contributes to the mesothelium of most organs of the coelomic cavity, including the heart, lungs, spleen, liver, stomach, and the intestine, being also essential for proper development and homeostasis of the kidneys and the urogenital system (24). During cardiogenesis, Wt1 contributes to epicardial progenitors that give rise to coronary vasculature and interstitial fibroblasts (FBs) through a process of epithelial to mesenchymal transition (25-27). Furthermore, Wt1 is expressed in a small fraction of cardiomyocytes (28, 29) and it has been reported that Wt1 is also expressed in postnatal non-coronary endothelial cells of the microvasculature (30). In the lungs, Wt1 contributes to the pulmonary mesenchyme forming the vascular and bronchial smooth muscle cells (SMCs), tracheal cartilage and part of the arterial endothelium, as well as to FB-like cells from the airways (31). Nevertheless, the adult lineage tracing of Wt1 in the lungs has not been evaluated to the best of our knowledge.

Here, we characterize a novel mouse model of HIF2 deletion in cardiopulmonary vascular and interstitial populations in response to chronic hypoxia using the *Wt1Cre* line (32). This model allows us to evaluate the impact of simultaneous elimination of HIF2 in ECs, PCs and SMCs, and hence is a valuable tool to anticipate potential effects of systemic HIF2 abrogation. Our data reveal that lack of HIF2 signaling in the Wt1 lineage protects against the elevation of the RVSP by preventing arteriolar muscularization, while it results in lung capillary leakage, alveolar hemorrhages, inflammation and pulmonary congestion. Moreover, elimination of HIF2 in the Wt1 compartment has detrimental effects on the cardiac adaptation to sustained low oxygen, leading to cardiomegaly, ventricular hypertrophy, dilatation and systolic dysfunction, together with microvasculature instability. Interestingly, most of these cardiopulmonary structural abnormalities are rescued after 1 week of reoxygenation, while cardiac function remains affected. Our data uncover novel roles of HIF2 in the cardiopulmonary system beyond the arterial endothelium in response to chronic hypoxia, and suggest that heart/lung HIF2 signaling exerts a positive function protecting against excessive capillary remodeling in response to low oxygen. Our work expands our knowledge on the role of HIF2 in the cardiovascular system during sustained hypoxia and might be relevant in the setting of novel pharmacological strategies based on HIF2 inhibition for PAH (33, 34).

## Results

### Wt1 lineage contributes to the macro and micro vascular compartments of heart and lungs

To further investigate Wt1 lineage in the adult heart, we performed lineage tracing analysis using a Rosa-tdTomato reporter mice. We confirmed the previously reported contribution of Wt1 lineage to coronary arteries (27, 35) including ECs [ETS related gene (ERG)^+^] and VSMCs [smooth muscle actin (SMA)^+^] (Figure 1A). Additionally, we identified that Wt1 lineage also contributes to PCs [Chondroitin Sulfate Proteoglycan 4 (CSPG or NG2)^+^] and interstitial FBs [podoplanin, (PDPN or GP38)+] surrounding and connecting the capillaries of the microvasculature (Figure 1B). We further investigated Wt1 lineage contribution to major cardiac cell populations by flow cytometry using the Rosa-tdTomato/Wt1Cre mice. FACS analysis showed that Wt1 lineage contributed to 31,4% of CD45^-^-non-myocyte cells in the adult heart (Figure 1C). Within the Tomato^+^-Wt1-derived cells, around 6.4% corresponds to CD39^+^/CD90^+^ VSMCs (Figure 1D), 24% to CD90^+^/GP38^-^ PCs (Figure 1E), 50.1% to CD31^+^ ECs (Figure 1F) and 8.4% to CD31^-^/CD90^-^/GP38^+^ FBs (Figure 1F, Table 1A). Furthermore, we calculated the relative contribution of Wt1 lineage within each non-myocyte population (Table 1B) and determined that there is a significant percentage of cardiac vascular cells contributed by the Wt1 lineage.

**Figure 1.**
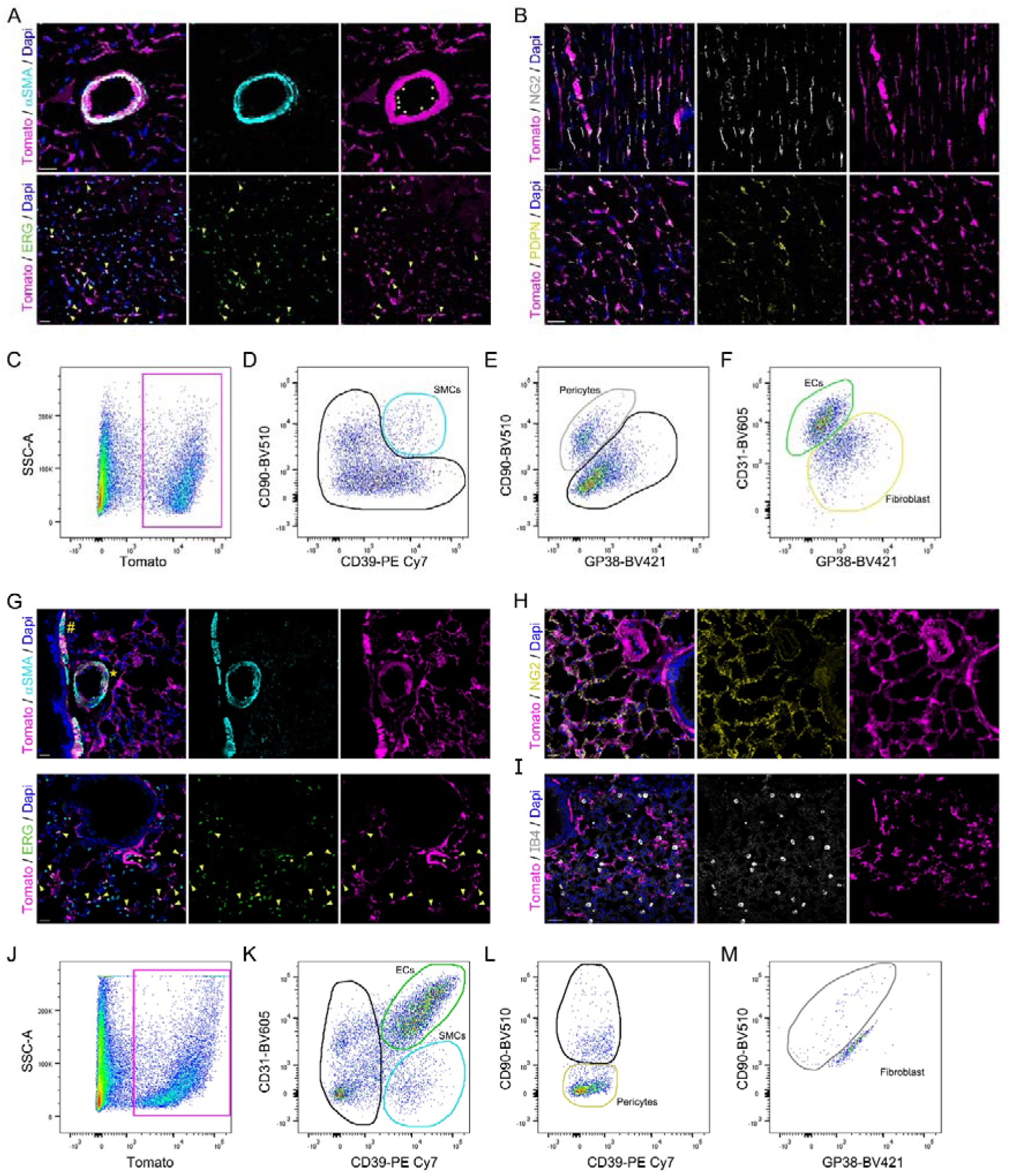
Wt1 lineage contribution to the heart and lung. **(A-B)** Immunofluorescence of cardiac sections from Rosa-tdTomato/Wt1Cre reporter mice with cell lineage markers. **A)** αSMA (VSMCs, cyan); ERG (ECs, green); Dapi (Nucleus, blue) and Tomato (Wt1 lineage, magenta). Upper panels show complete colocalization of αSMA^+^/Tomato^+^ signal and some ECs (yellow asterisks) in a coronary artery and lower panels reveal ERG^+^/Tomato^+^ ECs of capillaries (yellow arrowheads). Scale bars 20μm. **B)** Top panels show colocalization between NG2 (PCs, white) and Tomato (Wt1 lineage, magenta). Bottom panels are stained with PDPN (FBs, yellow) and Tomato (Wt1 lineage, magenta). Nucleus are co-stained with Dapi (blue). Scale bars 20μm. **(C-F)** Representative flow cytometry plots for identification of non-myocyte cardiac populations within the Wt1 lineage. **C)** Magenta rectangle contains Tomato^+^ cells within all CD45^-^ cells. **D)** Within Tomato^+^ cells in C, the cyan gate contains CD90^+^/CD39^+^ VSMCs. **E)** From the black gate on D, the gray gate corresponds to CD90^+^/GP38^-^ PCs. **F)** From PCs^-^ cells in E, the green gate represents CD31^+^/GP38^-^ ECs and the yellow gate CD31^-^/GP38^+^ FBs. **(G-I)** Immunofluorescence of lung sections of Rosa-tdTomato/Wt1Cre reporter mice with cell lineage markers. **G)** αSMA (VSMCs, cyan), ERG (ECs, green), Dapi (Nucleus, blue) and Tomato (Wt1 lineage, magenta). Upper panels show colocalization of αSMA^+^/Tomato^+^ in the bronchial submucosa (yellow hash) and in the medial layer of a pulmonary artery (yellow asterisk). Lower panels display alveolar ERG^+^/Tomato^+^ ECs (yellow arrowheads). Scale bars 20μm. **H, I)** Colocalization of Tomato (Wt1 lineage, magenta) with NG2 (**H**, PCs, yellow), but not with IB4, (**I**, alveolar macrophages, white). Nucleus are co-stained with Dapi (blue). Scale bars 20μm (**H**) and 40μm (**I**). **(J-M)** Representative plots of the flow cytometry strategy to characterize Wt1 pulmonary lineage. **J)** Magenta rectangle contains positive selection of Tomato^+^ cells within all CD45^-^ cells. **K)** Within Tomato^+^ cells, the cyan gate represents CD90^+^/CD31^-^ VSMCs and the green gate, CD31^+^/CD39^+^ ECs. **L)** From the VSMC-/EC- black gate in **K**, PCs were identified as CD39^-^/CD90^-^. **M)** Finally, we excluded cell autofluorescence detected with the single markers (data not shown) and kept the CD90^+^/GP38^-^as the FBs gate (grey).

**Table 1A.** Table of individual values and the mean and standard deviation (SD) from 5 independent experiments showing the percentage of each indicated cell type (ECs, PCs, FBs, SMCs) within the total Wt1 contribution to non-myocyte lineage in the heart.

| **% of each population within the Tomato<sup>+</sup>-Wt1 cardiac-lineage** |  |  |  |  |  |  |  |
| --- | --- | --- | --- | --- | --- | --- | --- |
| **Heart** |  |  |  |  |  |  |  |
| **Cell Type** | **Sample 1** | **Sample 2** | **Sample 3** | **Sample 4** | **Sample 5** | **Mean** | **SD** |
| **ECs** | 45,97 | 54,46 | 42,51 | 54,18 | 53,22 | **50,07** | 5,48 |
| **PCS** | 16,58 | 26,25 | 33,08 | 24,19 | 20,12 | **24,04** | 6,28 |
| **FBs** | 19,57 | 2,90 | 4,76 | 4,86 | 8,20 | **8,06** | 6,72 |
| **SMCs** | 3,13 | 7,42 | 7,87 | 6,42 | 7,18 | **6,40** | 1,90 |

**Table 1B.** Table of individual values and the mean and standard deviation (SD) from 5 independent experiments showing the percentage of contribution of Wt1-derived cells to each cell lineage in the heart.

| **% of Wt1 lineage contribution within each non-myocyte population** |  |  |  |  |  |  |  |
| --- | --- | --- | --- | --- | --- | --- | --- |
| **Heart** |  |  |  |  |  |  |  |
| **Cell Type** | **Sample 1** | **Sample 2** | **Sample 3** | **Sample 4** | **Sample 5** | **Mean** | **SD** |
| **ECs** | 14,59 | 25,71 | 21,75 | 28,76 | 24,26 | **23,01** | 5,35 |
| **PCS** | 76,13 | 83,60 | 86,44 | 79,74 | 80,52 | **81,29** | 3,92 |
| **FBs** | 75,37 | 41,67 | 60,77 | 46,92 | 57,50 | **56,44** | 13,11 |
| **SMCs** | 75,64 | 78,42 | 58,55 | 75,08 | 78,93 | **73,32** | 8,43 |

Regarding the adult lung, using the same reporter model of Rosa-tdTomato/Wt1Cre, we found that Wt1 lineage contributes to ECs of the alveolar capillary network, small and large arteries, SMCs in the medial layer of pulmonary arteries and in the bronchial submucosa (Figure 1G), as well as to PCs (Figure 1H). In contrast, there is no contribution of Wt1 lineage to alveolar type I (AT1) or type II (AT2) cells identified by Retinoic Acid Inducible protein 3 (Rai3), also known as G Protein-Coupled Receptor Class C Group 5 Member A (GPRC5A) (36) or Surfactant Protein-C (SPC) (37, 38) respectively (Figure S1A), neither to alveolar macrophages labelled by isolectin B4 (IB4) (39) (Figure 1I). As in the heart, we further investigated Wt1 lineage contribution to major lung cell populations by flow cytometry using the Rosa-tdTomato/Wt1Cre mice. FACS analysis showed that over 18.3% of the CD45^-^ fraction of adult lung expresses or is Wt1-derived (Figure 1J). Out of these Tomato^+^ cells, around 37.3% corresponds to CD31^+^/CD39^+^ endothelium (Figure 1K), 20.6% to CD31^-^/CD39^+^ SMCs (Figure 1K), 22.8% CD90^-^/GP38^-^ to PCs (Figure 1L), as further confirmed by a NG2-DsRed reporter mice (Figure S1B), and 1.7% CD90^+^ FBs (Figure 1L, M, Table 2A). The relative contribution of Wt1-derived cells to pulmonary lineages was also determined (Table 2B).

**Table 2A.** Table of individual values and the mean and standard deviation (SD) from 5 independent experiments showing the percentage of each indicated cell type (ECs, PCs, FBs, SMCs) within the total Wt1 contribution in the lung.

| **% of each population within the Tomato<sup>+</sup>-Wt1 lung lineage** |  |  |  |  |  |  |  |
| --- | --- | --- | --- | --- | --- | --- | --- |
| **Lung** |  |  |  |  |  |  |  |
| **Cell Type** | **Sample 1** | **Sample 2** | **Sample 3** | **Sample 4** | **Sample 5** | **Mean** | **SD** |
| **ECs** | 54,93 | 35,41 | 26,83 | 31,48 | 37,70 | **37,27** | 10,70 |
| **PCS** | 16,08 | 25,86 | 27,90 | 21,05 | 23,07 | **22,79** | 4,57 |
| **FBs** | 1,23 | 1,23 | 1,49 | 1,18 | 1,68 | **1,36** | 0,22 |
| **SMCs** | 6,50 | 19,05 | 26,13 | 33,40 | 17,72 | **20,56** | 10,05 |

**Table 2B.** Table of individual values and the mean and standard deviation (SD) from 5 independent experiments showing the percentage of contribution of Wt1-derived cells to each cell lineage in the lung.

| % of Wt1 contribution within each cell population |  |  |  |  |  |  |  |
| --- | --- | --- | --- | --- | --- | --- | --- |
| Lung |  |  |  |  |  |  |  |
| Cell Type | Sample 1 | Sample 2 | Sample 3 | Sample 4 | Sample 5 | Mean | SD |
| **ECs** | 22,67 | 14,03 | 13,33 | 36,22 | 16,98 | **20,65** | 9,45 |
| **PCS** | 14,69 | 9,62 | 10,25 | 22,00 | 11,54 | **13,62** | 5,08 |
| **FBs** | 5,15 | 2,32 | 1,90 | 4,78 | 5,11 | **3,85** | 1,60 |
| **SMCs** | 18,61 | 16,02 | 18,81 | 36,70 | 12,61 | **20,55** | 9,37 |

In summary, these lineage tracing analyses demonstrated that Wt1 lineage contributes to macro and microvasculature of heart and lung, offering an important tool for genetic manipulation and molecular analysis of the different cell components involved in vascular function.

### Elimination of Hif2 in the Wt1 lineage prevents pulmonary arteriole muscularization and protects against elevation of the RVSP upon chronic hypoxia

In order to understand the role of HIF2 in the cardiovascular and pulmonary compartments contributed by Wt1 (Figure 1), we generated a new conditional HIF2 knock out model by crossing the HIF2-floxed (40) line with the *Wt1Cre* mouse line (32), from now on *Hif2/Wt1* cKO. First, we confirmed efficient deletion of *Hif2* floxed exon 2 by PCR (data not shown). Next, we determined that elimination of *Hif2* in the Wt1 lineage does not cause embryonic lethality or any obvious cardiovascular structural defect, including the proper formation of the ventricular chambers or the coronary tree (Figure S2A). Furthermore, we confirmed that *Hif2/Wt1* cKO display normal survival curve from weaning to adulthood (Figure S2B), suggesting that Wt1/HIF2 signaling is not required for the correct formation and homeostasis of the heart. Afterwards, we evaluated lung tissue integrity under normoxic conditions on whole mount (not shown) and by hematoxylin and eosin (HE) staining of the lung parenchyma (Figure 2A), finding no obvious differences between the structure of control and *Hif2/Wt1* cKO mice in basal conditions. Thereafter, we investigated the impact of *Hif2* deletion in the Wt1 lineage in response to chronic hypoxia. To that aim, we exposed 12 weeks old control and *Hif2/Wt1* cKO mice to 10% oxygen during 2 or 3 weeks and evaluated functional and structural parameters by echography and classical histology (Figure S3). Because it has been previously reported that hypoxia induces vascular remodeling by muscularization of distal pulmonary arterioles (17, 41), first we performed an SMA staining in lung sections in normoxia or after exposure to hypoxia (Figure 2B). Tissue analysis revealed that after 2 and 3 weeks of hypoxia, there were no significant differences between control and mutant mice on the number of large (30-20 µm) (Figure 2C) neither on medium (19-10 µm) caliber arterioles (Figure 2D), although there was an upward tendency in the number of medium arterioles of control mice respect to *HIF2/Wt1* cKO mice. This difference became highly significant for the number of small arteries (9-1 µm) found in control mice after 2 and 3 weeks of hypoxia compared to normoxic conditions, while *Hif2/Wt1* cKO mice exhibited a similar number of small arteries in both conditions (Figure 2E). Then, we determined the effect of the vascular remodeling differences after hypoxia exposure in the measurement of the RVSP. As expected, control mice displayed a significant elevation of the RVSP after 2 and 3 weeks of chronic hypoxia. In contrast, the *Hif2/Wt1* cKO mice were protected against the elevation of RVSP at both time points, showing similar pressures in normoxia or hypoxia conditions (Figure 2F). These results are in agreement with former works describing the role of endothelial HIF2 in arterial remodeling upon chronic hypoxia (18-20).

**Figure 2.**
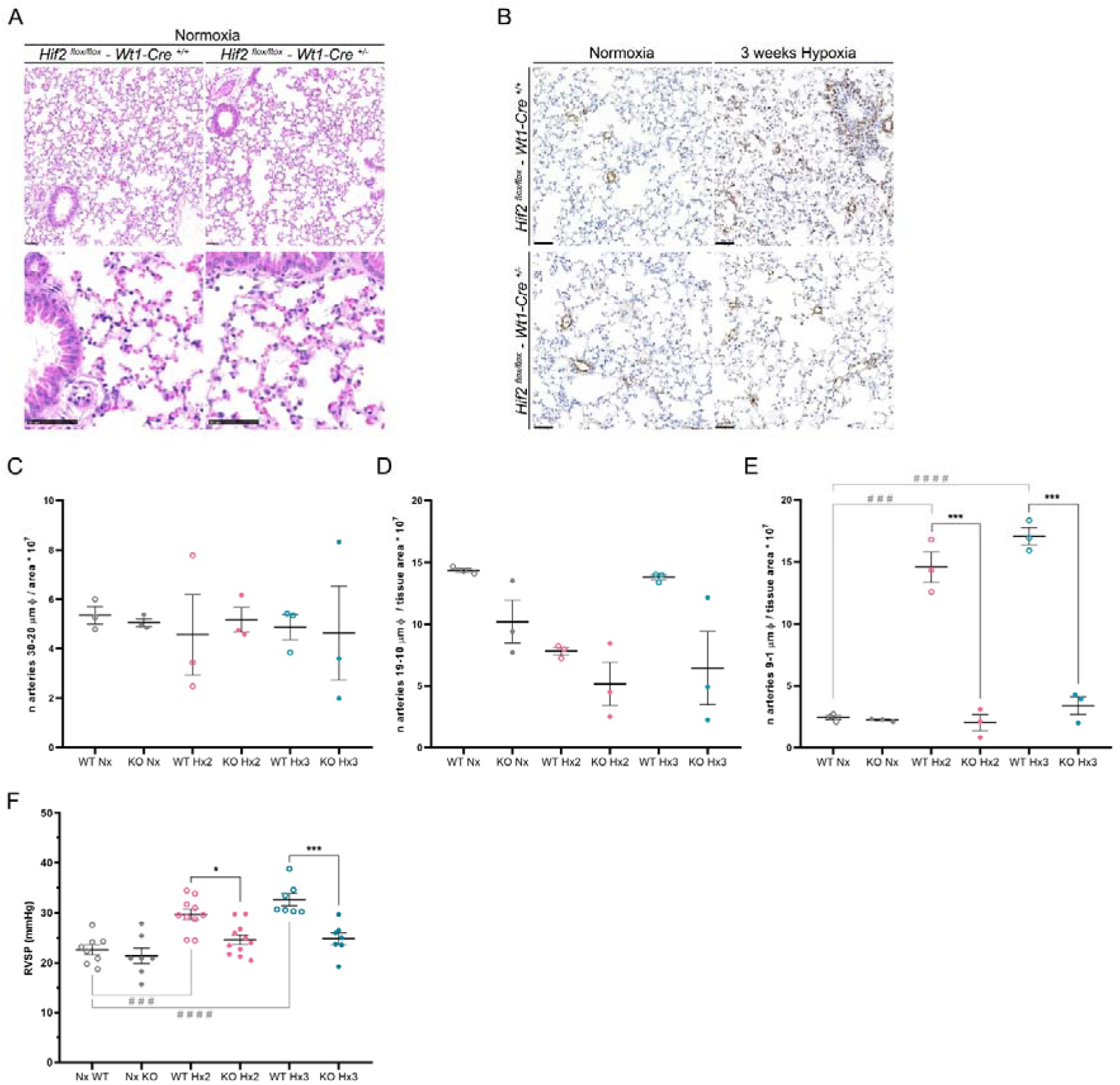
Muscularization of pulmonary arteries under chronic hypoxia. **A)** Histological analysis by HE of lung sections from control (Hif2^flox/flox^-Wt1-Cre^+/+^, left panels) and *Hif2/Wt1* cKO mice (Hif2^flox/flox^-Wt1-Cre^+/-^, right panels) in normoxic conditions. Scale bars 50μm. **B)** Representative images of αSMA immunohistochemistry in control (Hif2^flox/flox^-Wt1-Cre^+/+^, top panels) and *Hif2/Wt1* cKO (Hif2^flox/flox^-Wt1-Cre^+/-^, bottom panels) mice in normoxia or chronic (3 weeks) hypoxia. Scale bar 50μm. **(C-E)** Quantification of the number of lung arterioles ranging from 30-20μm (**C**), 19-10μm (**D**) or 9-1μm (**E**) per tissue area in control (WT) and *Hif2/Wt1* cKO (KO) mice in normoxia (Nx, gray dots) of after 2 (Hx2, pink dots) and 3 (Hx3, blue dots) weeks of sustained hypoxia. **F)** Scatter dot plot of the right ventricular systolic pressure (RVSP). All graph bars show individual values (dots) and the black line represents the Mean ± SEM. Anova test, p value: * P ≤ 0,05; ** P ≤ 0,01; *** P ≤ 0,001; **** P ≤ 0,0001.

Altogether these data indicate that elimination of HIF2 in the Wt1 lineage prevents vascular remodeling and subsequent RVSP elevation after chronic hypoxia, protecting against hallmarks of PAH in response to low oxygen.

### Functional HIF2 in the microvascular compartment is necessary for alveolar parenchyma stability and pulmonary performance during sustained hypoxia

To further investigate the importance of HIF2 signaling in the pulmonary Wt1 lineage, we performed lung and right-sided cardiac echography analysis in control and *Hif2/Wt1* cKO mice in normoxia and after 2 or 3 weeks of chronic hypoxia at 10% O_2_. First, we determined the value of the pulmonary artery acceleration time *versus* ejection time ratio (AT/ET) indicative of the pulmonary artery pressure, finding no significant differences between control and *Hif2/Wt1* cKO mice neither in normoxia, nor after 2 or 3 weeks of hypoxia (Figure 3A). Next, we calculated the mouse lung ultrasound score (MoLUS score) that predicts the level of pulmonary congestion integrating several functional and structural parameters of the lung (pleural effusion, alveolar edema/hemorrhages, presence or absent of A, B and Z lines among others), and that has been previously reported to correlate with cardiac function (42). Echographic analysis confirmed similar MoLUS values between control and *Hif2/Wt1* cKO mice in normoxia and after 2 weeks of hypoxia exposure, but a notable increase on this parameter in Hif2 mutants after 3 weeks of sustained hypoxia (Figure 3B, 3C). These results indicated that, despite their protection against vascular remodeling (Figure 2C) and RVSP elevation (Figure 2G), *Hif2/Wt1* cKO mice displayed worse pulmonary performance and profound structural abnormalities during sustained hypoxia. To further investigate the extent of lung congestion in *Hif2/Wt1* cKO upon hypoxia, we analyzed the structure of the pulmonary parenchyma by HE staining, observing an important thickening of the alveolar wall in the *Hif2/Wt1* cKO mice relative to controls by 2 weeks of hypoxia (Figure 3D). Moreover, by 3 weeks of hypoxia the *Hif2/Wt1* cKO mice lung displayed severe erythrocyte congestion in the alveolar parenchyma, arterioles and arteries (Figure 3D). This increased alveolar wall thickening, and hemorrhages resulted in a significant reduction of the alveolar space in *Hif2/Wt1* cKO mice compared to controls after both 2 and 3 weeks of hypoxia exposure (Figure 3E). In addition, after 3 weeks of chronic hypoxia, there was an increased amount of alveolar macrophages, many of them loaded with erythrocytes as evidenced by positive hemosiderin signal on HE staining (Figure 3F). We further confirmed the increase in the number of alveolar macrophages in the *Hif2/Wt1* cKO lungs by 3 weeks of hypoxia by immunofluorescence with IB4 (Figure 3G, 3H) and its expansion with the proliferation marker Ki67 (Figure 3G, Figure 3I). Since Wt1 lineage does not contribute neither to alveolar macrophages (Figure 1I) nor to AT1 or AT2 cells (Figure S1A), our results suggest that these alveolar alterations in the *Hif2/Wt1* cKO mice might be indirect and secondary to the altered vascular remodeling occurring in HIF2 mutants in response to sustained hypoxia. To evaluate this hypothesis, we analyzed whether the endothelium of the alveolar parenchyma was affected in the *Hif2/Wt1* cKO. Immunostaining with markers for ECs (ERG) and proliferation (Ki67) after exposure to chronic hypoxia (Figure 3G), showed a significant increase in the number of proliferating ECs in the *Hif2/Wt1* cKO mice compared to controls (Figure 3J).

**Figure 3.**
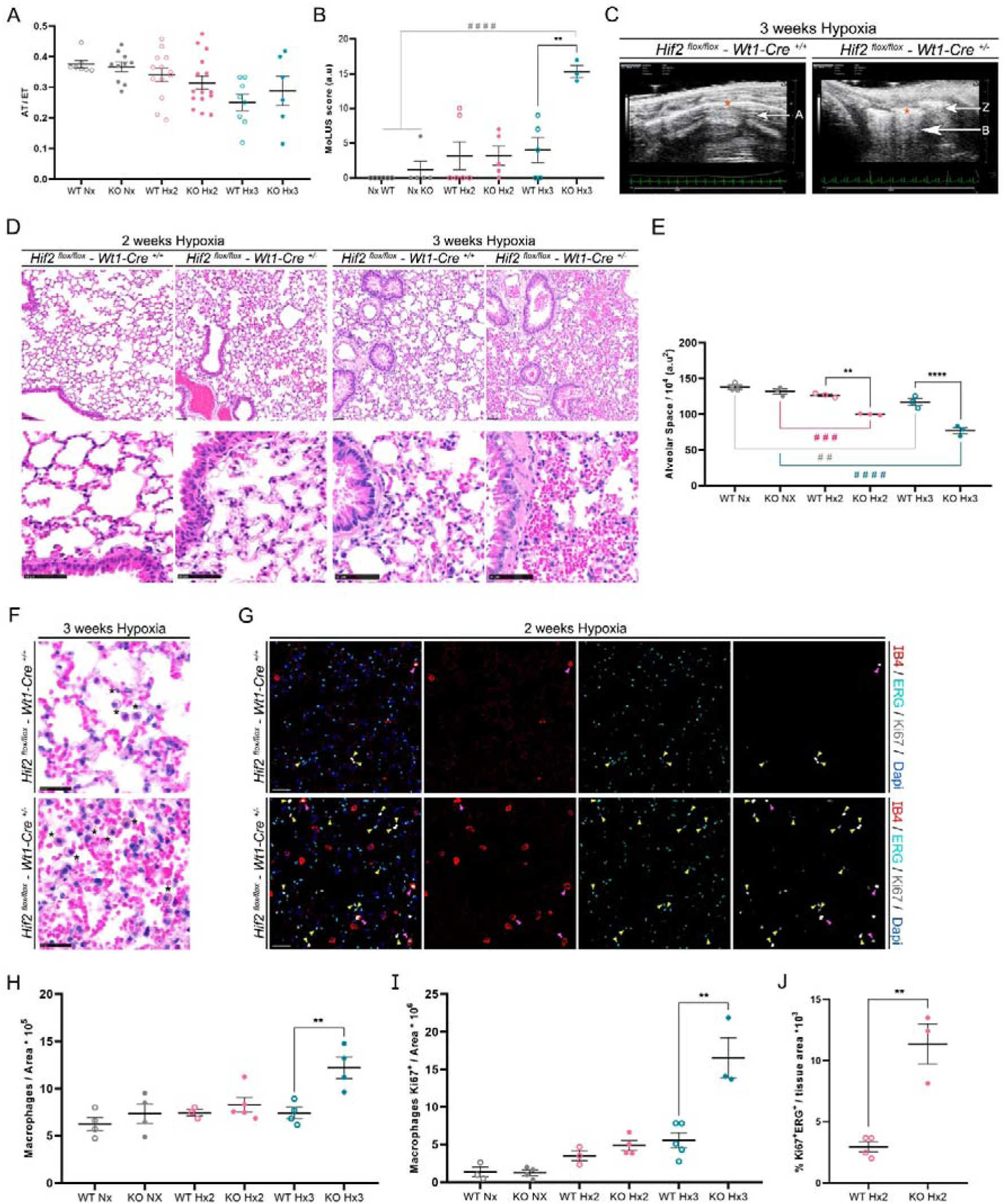
Pulmonary architecture and function during sustained low oxygen conditions. **A)** Scatter dot plot of pulmonary artery acceleration to ejection time ratio (AT/ET) measured by Doppler echography in control (WT) and *Hif2/Wt1* cKO (KO) mice in Normoxia (Nx, gray dots), 2 weeks hypoxia (Hx2, pink dots) or 3 weeks hypoxia (Hx3, blue dots). **B)** Scatter dot plot of Mouse Lung ultrasound score (MoLUS) of control (WT) and *Hif2/Wt1* cKO (KO) mice in normoxia (Nx gray dots) and hypoxia conditions [2 (Hx2, pink dots) or 3 weeks hypoxia (Hx3, blue dots)]. **C)** Representative lung echography images of control (Hif2^flox/flox^-Wt1-Cre^+/+^, left panel) and *Hif2/Wt1* cKO (Hif2^flox/flox^-Wt1-Cre^+/-^, right panel) mice after 3 weeks of hypoxia exposure. B (white large tracks from top to bottom) and Z lines (white short lines under the pleura) reflecting pulmonary edema are obvious in the *Hif2/Wt1* cKO compared to controls with only A lines (fine lines parallel to the pleural line). Pleural thickening and fragmentation are evident in *Hif2/Wt1* cKO (white intense line, orange asterisk on right panel) compared to normal pleura indicative of well-aerated lung (white thin line, orange asterisk on left panel. **D)** Histopathological characterization by HE staining of lung sections of control (Hif2^flox/flox^-Wt1-Cre^+/+^, left panels of each time of hypoxia) and *Hif2/Wt1* cKO (Hif2^flox/flox^-Wt1-Cre^+/-^, right panels of each time of hypoxia) mice. Scale bars 50μm. **E)** Scatter dot plot showing the quantification of the alveolar space in control (WT) and *Hif2/Wt1* cKO (KO) mice in normoxia (Nx, gray dots) and hypoxic conditions [2 (Hx2, pink dots) or 3 weeks hypoxia (Hx3, blue dots)]. **F)** Representative HE images of pulmonary inflammation showing alveolar macrophages (black asterisks) in control (Hif2^flox/flox^-Wt1-Cre^+/+^, top panel) and *Hif2/Wt1* cKO (Hif2^flox/flox^-Wt1-Cre^+/-^, bottom panel) mice after 3 weeks of hypoxia. Scale bars 25μm. **G)** Representative immunofluorescence of pulmonary sections from control (Hif2^flox/flox^-Wt1-Cre^+/+^, top panels) and *Hif2/Wt1* cKO (Hif2^flox/flox^-Wt1-Cre^+/-^, bottom panels) stained for ERG (ECs, cyan); Ki67 (mitosis, white); IB4 (alveolar macrophages, red) and Dapi (nucleus, blue) showing proliferation of ECs (ERG^+^/Ki67^+^, yellow arrowheads) and alveolar macrophages (IB4^+^/Ki67^+^, pink arrowheads) after 2 weeks of Hx. Scale bars 20μm. **H)** Scatter dot plot showing the quantification of macrophages per tissue area in controls (WT) and *Hif2/Wt1* cKO (KO) mice in normoxia (Nx, gray dots) and hypoxic conditions [2 (Hx2, pink dots) or 3 weeks hypoxia (Hx3, blue dots)]. **I)** Number of proliferating macrophages per tissue area in lung sections from control (WT) and *Hif2/Wt1* cKO (KO) mice in normoxia (Nx, gray dots) and hypoxia conditions [2 (Hx2, pink dots) or 3 weeks hypoxia (Hx3, blue dots)]. **J)** Percentage of ECs proliferation (ERG^+^/Ki67^+^) per tissue area in control (WT) and *Hif2/Wt1* cKO (KO) mice after 2 weeks of Hx exposure (Hx2, pink dots). All graph bars show individual values (dots) and the black line represents the Mean ± SEM. Anova test, p value: * P ≤ 0,05; ** P ≤ 0,01; *** P ≤ 0,001; **** P ≤ 0,0001.

Finally, FACS analysis from lung tissue of Rosa-tdTomato/Wt1Cre reporter control or HIF2 floxed-Rosa-tdTomato/Wt1Cre (*Hif2/Wt1* reporter cKO) mice revealed a significant elevation in total cells contributed by the Wt1 lineage (Tomato^+^) in *Hif2/Wt1* reporter cKO compared with reporter control mice after 2 weeks of 10% O_2_ exposure or with *Hif2/Wt1* reporter cKO mice in normoxia (Figure 4A). These quantitative changes by FACS analysis were further confirmed qualitatively by immunofluorescence (Figure 4B). The increase in total Tomato^+^ cells could be explained by the elevation on Tomato^+^-ECs in the *Hif2/Wt1* reporter cKO in hypoxia relative to normoxia and compared with reporter control mice after 2 weeks of hypoxia (Figure 4C, 4D). In contrast, we observed no significant changes in the total Tomato^+^-PCs in the lung between normoxia or hypoxia in reporter control, neither in *Hif2/Wt1* reporter cKO mice, although there is a trend of induction in Tomato^+^-PCs in the *Hif2/Wt1* reporter cKO compared to control mice after 2 weeks of chronic hypoxia (Figure 4E, 4F).

**Figure 4.**
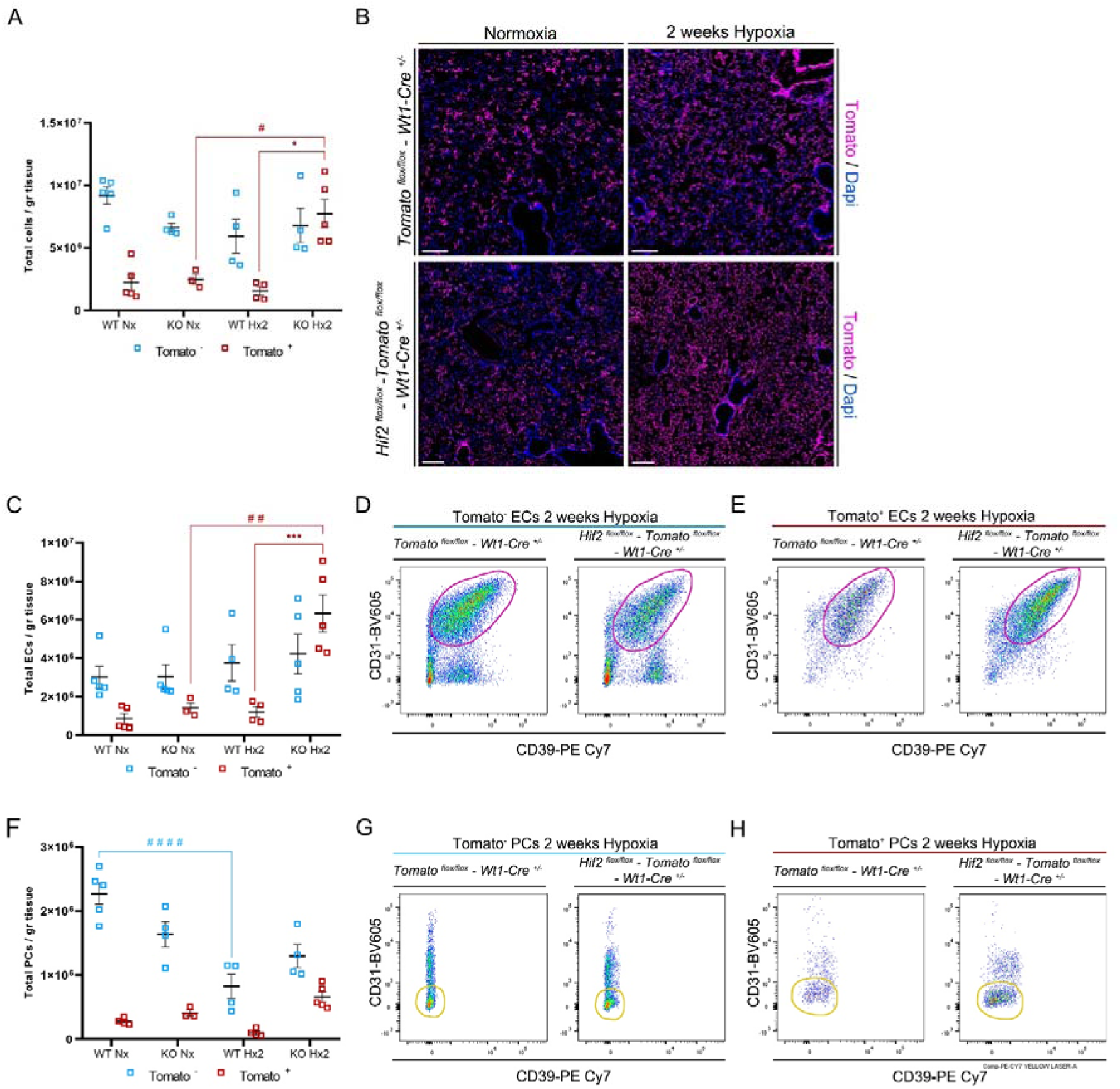
Quantitative analysis of Wt1-derived pulmonary cells during chronic hypoxia. **A)** Quantification of total cells per gram of tissue of Tomato^-^ (blue squares) and Tomato^+^ (red squares) cells in Rosa-tdTomato/Wt1 reporter control (WT) and Rosa-tdTomato/Hif2 reporter cKO(KO) mice in normoxia (Nx) and after 2 weeks of hypoxia (Hx2) analyzed by FACS. **B)** Immunofluorescence of Tomato (Wt1 lineage, magenta) and Dapi (nucleus, blue) in lung sections from Rosa-tdTomato/Wt1 reporter control (Tomato^flox/flox^-Wt1^+/-^, top panels) and Rosa-tdTomato/Hif2 reporter cKO (Tomato^flox/flox^/Hif2^flox/flox^/Wt1^+/-^), bottom panels) mice in normoxia (left column) and after 2 weeks of hypoxia (right column). Scale bars 200μm. **C, E)** Scatter dot plot representing changes in the total cells per gram of tissue of Tomato^-^ (blue squares) and Tomato^+^ (red squares) ECs (**C**) or PCs (**E**) in control (WT) and mutant (KO) mice in normoxia (Nx) and after 2 weeks of hypoxia (Hx2). **D, E, G, H)** Representative FACS pseudocolor image showing Tomato^-^-ECs (purple gates, **D**) and Tomato^+^-ECs (purple gates, **E**) or Tomato^-^-PCs (yellow gates, **G**) and Tomato^+^-PCs (yellow gates, **G**) from control (Tomato^flox/flox^-Wt1^+/-^) and mutant (Tomato^flox/flox^-Hif2^flox/flox^-Wt1^+/-^) lung tissue. All panels show the ECs and PCs content from a fixed number of 24000 CD45^-^ cells. For all scatter plots individual values are shown and the black line represents the Mean ± SEM. Anova test. P: pvalue. * P ≤ 0,05; ** P ≤ 0,01; *** P ≤ 0,001; **** P ≤ 0,0001.

In summary these results reveal that despite of the protective effect on preventing pulmonary arteriole remodeling and elevation of RVSP, defective HIF2 signaling in the Wt1 compartment is deleterious for lung adaptation to chronic hypoxia. Indeed, HIF2 deletion in the Wt1 lineage results in unstable and proliferative microvasculature that favors erythrocyte extravasation and macrophage proliferation, leading to alveolar wall thickening, severe lung congestion and pulmonary damage in response to chronic hypoxia.

### Hif2/Wt1 cKO mutants display cardiac hypertrophy and ventricular dilation in response to low oxygen exposure

Since the Wt1 lineage has a broad contribution to several vascular compartments of the heart (Figure 1A-B) and considering that the direct role of HIF2 in cardiac function and tissue structure in response to chronic hypoxia remains elusive, we decided to explore the impact of HIF2 deletion in the Wt1 cardiac lineage upon sustained low oxygen exposure. To this aim, we exposed 12 weeks old control and *Hif2/Wt1* mutant mice to 10% O_2_ during 2 and 3 weeks (Figure S3) and performed cardiac functional and structural analysis before and after hypoxia treatment. Echocardiography characterization revealed that the lack of HIF2 in the Wt1 lineage favors cardiac hypertrophy of both, the right (RV), and specially the left ventricle (LV) in response to low oxygen tensions, while controls remained unaltered (Figure 5A, 5B). Whereas right ventricular systolic function (TAPSE) was unaffected in mutant mice relative to controls after exposure to chronic hypoxia (Figure 5C), *Hif2/Wt1* cKO mice displayed common hallmarks of left ventricular systolic heart failure, with a significant reduced LV ejection fraction (LVEF, Figure 5D) and increased end diastolic volume (LVED Vol, Figure 5E), indicative of cardiac dilatation (Figure 5F). In order to connect functional parameters with structural adaptations, next we performed whole mount organ analysis, finding a progressive increase of the heart weight to body weight ratio in *Hif2/Wt1* cKO mutant mice relative to controls (Figure 5G). This gradual cardiomegaly in response to chronic hypoxia was further validated by HE staining (Figure 5H).

**Figure 5.**
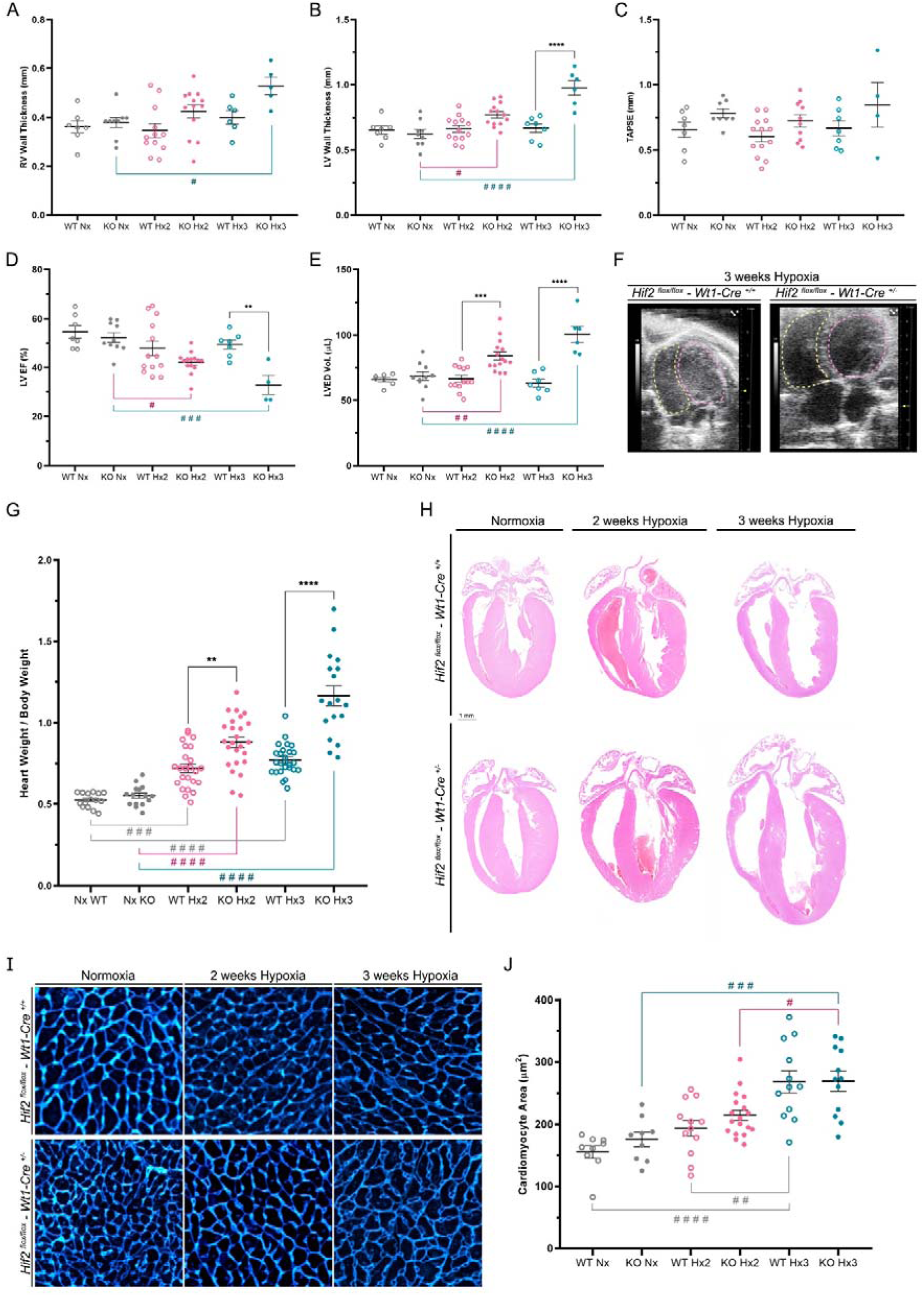
Cardiac functional and structural characterization in chronic hypoxia. **(A-E)** Functional analysis of the heart by echocardiography of control (WT) and *Hif2/Wt1* cKO (KO) mice in normoxia (Nx, grey dots), 2 (Hx2, pink dots) or 3 weeks hypoxia (Hx3, blue dots). **A)** Right ventricular (RV) wall thickness. **B)** Left ventricular (LV) wall thickness. **C)** Tricuspid annular plane systolic excursion (TAPSE). **D)** Left ventricular ejection fraction (LVEF). **E)** Left ventricular end diastolic volume (LVED Vol). **F)** Representative images of a 2D view showing the apical-four chamber view in a control (Hif2^flox/flox^-Wt1-Cre^+/+^, left panel) and *Hif2/Wt1* cKO (Hif2^flox/flox^-Wt1-Cre^+/-^, right panel) after 3 weeks of hypoxia. The yellow dashed line delimits the inner face of the RV, while the pink dashed line defines the inner face of the LV. **G)** Quantification of the heart weight to body weight ratio in control (WT) and *Hif2/Wt1* cKO (KO) mice in normoxia (Nx, gray dots) and 2 (Hx2, pink dots) or 3 weeks of Hx (Hx3, blue dots). **H)** Representative histological analysis by HE staining of heart sections from control (Hif2^flox/flox^-Wt1-Cre^+/+^, top panels) and *Hif2/Wt1* cKO (Hif2^flox/flox^-Wt1-Cre^+/-^, bottom panels) mice in normoxia (left), 2 weeks (middle) or 3 weeks of hypoxia (right). Scale bar 1mm. **I)** Representative images of cardiac sections from control (Hif2^flox/flox^-Wt1-Cre^+/+^, top panels) and *Hif2/Wt1* cKO (Hif2^flox/flox^-Wt1-Cre^+/-^, bottom panels) Wheat Germ Agglutinin (WGA) staining mask in Fiji to analyze cardiomyocyte size in normoxia (left columns), 2 weeks (middle columns) and 3 weeks of hypoxia (right columns). **J)** Quantification of the cardiomyocyte area in controls (WT) and *Hif2/Wt1* cKO (KO) mice in in normoxia (Nx, gray dots) and 2 (Hx2, pink dots) or 3 weeks of Hx (Hx3, blue dots). For all scatter plots individual values are shown and the black line represents the Mean ± SEM. Anova test. P: pvalue. * P ≤ 0,05; ** P ≤ 0,01; *** P ≤ 0,001; **** P ≤ 0,0001.

Considering that Wt1 lineage contributes to some scattered cardiomyocytes (29) as we further confirmed by lineage tracing (data not shown), we decided to explore whether the absence of *Hif2* could influence cardiomyocyte size upon hypoxia. To this aim, we performed wheat germ agglutinin (WGA) staining in 14-15 weeks old heart sections from control and *Hif2/Wt1* cKO mice and compared the size of cardiomyocytes in normoxia and after 2 and 3 weeks of exposure to 10% O_2_ in both groups (Figure 5I). The analysis indicated an ongoing increase in the cardiomyocyte cell area from normoxia to 2 and 3 weeks of hypoxia in both, control and *Hif2/Wt1* mutants (Figure 5J). Nevertheless, we did not find significant differences between control and *Hif2/Wt1* cKO mice at any time point, excluding cardiomyocyte cell hypertrophy as the mechanism responsible for the increased cardiomegaly and elevated heart weight to body weight ratio developed by *Hif2/Wt1* mutants after hypoxia.

Overall, these results indicate that deletion of Hif2 in the Wt1 lineage compromises cardiac adaptation to chronic hypoxia, causing LV systolic dysfunction and enhanced cardiomegaly not due to cardiomyocyte hypertrophy, pointing to a protective role of vascular HIF2 signaling in the heart during adaptation to low oxygen conditions.

### HIF2 signaling represses cardiac EC proliferation and prevents microvascular remodeling during chronic hypoxia

Next, we decided to assess whether the increased cardiomegaly of the *Hif2/Wt1* cKO during chronic hypoxia could be due to enhanced cardiac hyperplasia. To explore this possibility, we determined the proliferation rate by immunostaining of cardiac sections of control and *Hif2/Wt1* mutants in normoxia or after exposure to 2 and 3 weeks of hypoxia using mitosis markers [phospho histone H3 (PH3) and Ki67] together with WGA to label cardiomyocytes contour and IB4 and ERG for ECs. Image analysis revealed that hypoxia exposure did not influence cardiomyocyte proliferation, but rather promoted mitosis of the microvasculature (Figure 6A). We further confirmed that IB4^+^ proliferating cells were indeed ERG^+^ ECs (Figure S4A). Interestingly, EC proliferation followed a dynamic pattern, with a peak after 2 weeks of hypoxia exposure, followed by a drop close to normoxic values after 3 weeks of hypoxia (Figure 6B), suggesting that HIF2 regulates an inhibitory feedback loop to compensate cardiac endothelial proliferation induced by hypoxia. To explore the impact of proliferation in response to hypoxia, we evaluated the integrity of the cardiac capillary network using IB4 to mark the outline of ECs. Our analysis revealed an increase in the capillary area in the *Hif2/Wt1* cKO mice after 2 and 3 weeks of hypoxia that was not observed in the controls, neither in normoxia conditions (Figure 6C, 6D). This enlargement in capillary area suggested a higher capillary diameter that was accompanied by a reduction in the capillary density assessed as number of capillaries per tissue area (Figure 6E).

**Figure 6.**
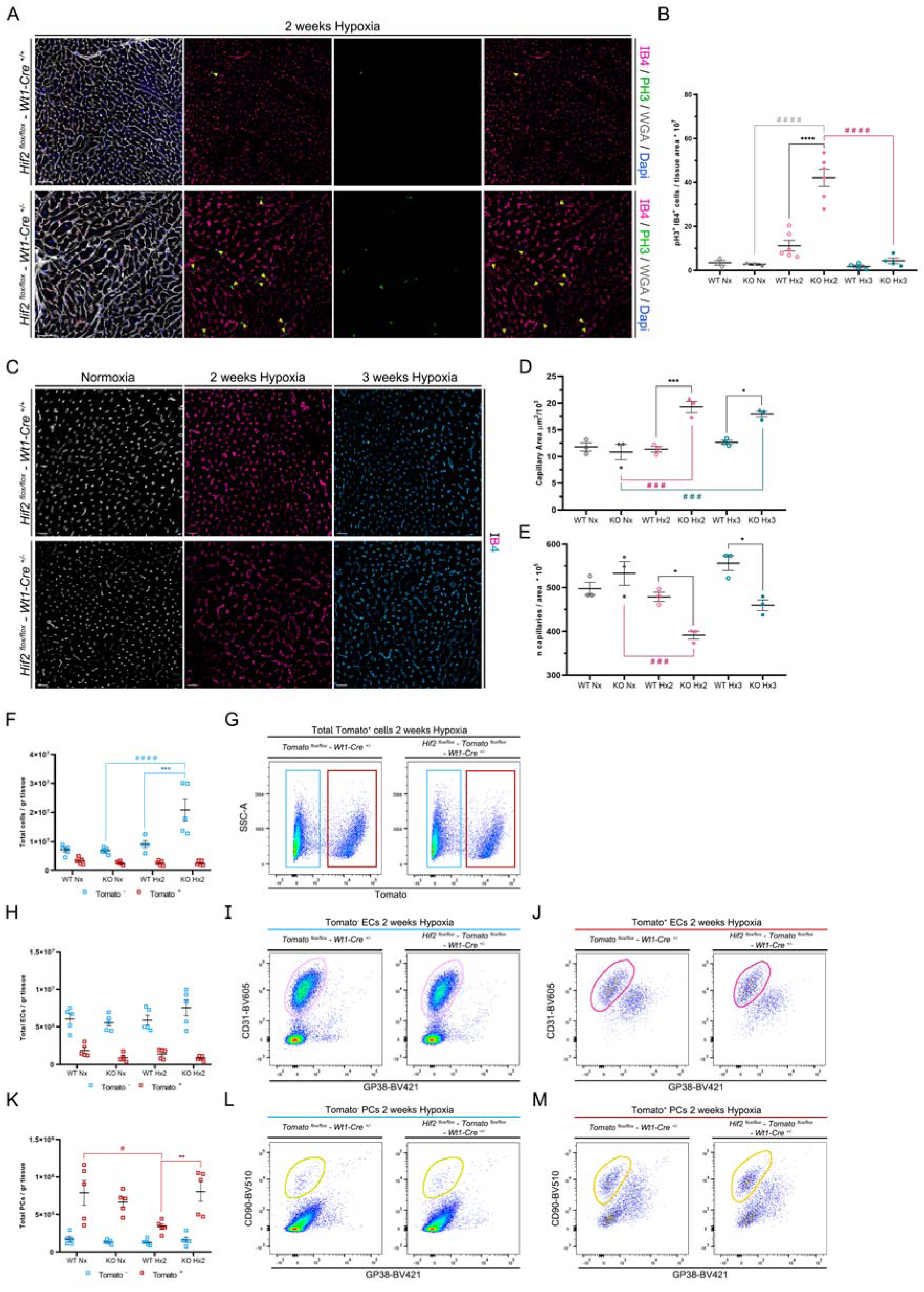
Cardiac microvascular remodeling in response to low oxygen. **A)** Immunofluorescence of representative cardiac sections after 2 weeks of hypoxia of control (Hif2^flox/flox^-Wt1-Cre^+/+^, top panels) and *Hif2/Wt1* cKO (Hif2^flox/flox^-Wt1-Cre^+/-^, bottom panels) mice with IB4 (outline of ECs, magenta), WGA (outline of all cells, white), PH3 (mitosis, green) and Dapi (nucleus, blue). Yellow arrowheads indicate IB4^+^/PH3^+^ cells. Scale bars 40μm. **B)** Quantification of the EC proliferation rate per tissue area in controls (WT) and *Hif2* mutants (KO) in normoxia (Nx, gray dots) and 2 (Hx2, pink dots) or 3 weeks of Hx (Hx3, blue dots). **C)** Representative images of IB4 immunofluorescence of control (Hif2^flox/flox^-Wt1-Cre^+/+^, top panels) and *Hif2/Wt1* cKO (Hif2^flox/flox^-Wt1-Cre^+/-^, bottom panels) mice in normoxia (left columns), 2 weeks (middle columns) and 3 weeks of hypoxia (right columns). Scale bars 20μm. **D, E)** Scatter dot plots for the quantification of the capillary area (**D**) and density (**E,** number of capillaries per tissue area) of controls (WT) and *Hif2* mutants (KO) in normoxia (Nx, gray dots) and 2 (Hx2, pink dots) or 3 weeks of Hx (Hx3, blue dots). **F-M.** FACS analysis with Rosa-tdTomato/Wt1 reporter control (WT) and Rosa-tdTomato/Hif2 reporter cKO (KO) mice in normoxia (Nx) and after 2 weeks of hypoxia (Hx2). **F, H, K)** Scatter dot plots showing Tomato^-^ (blue squares) and Tomato^+^ (red squares) total cardiac cells (**F**), ECs (**H**) and (PCs) (**K**) per gram (gr) of tissue. **G, I, J, L, M.** Representative FACS pseudocolor plots of control (Tomato^flox/flox^/Wt1^+/-^) and mutant (Tomato^flox/flox^/Hif2^flox/flox^/Wt1^+/-^) mice. All gates were generated from a fixed number of 34000 CD45^-^ cells. **G)** Total CD45^-^ cells separating Tomato^-^ (blue gate) and Tomato^+^ cells (red gate). Tomato^-^-ECs (purple gates, **I**) and Tomato^+^-ECs (purple gates, **J**) or Tomato^-^-PCs (yellow gates, **L**) and Tomato^+^-PCs (yellow gates, **M**). For all scatter plots individual values are shown and the black line represents the Mean ± SEM. Anova test. P: pvalue. * P ≤ 0,05; ** P ≤ 0,01; *** P ≤ 0,001; **** P ≤ 0,0001.

Surprisingly, and in contrast to our observations in the lung, FACS analysis using the Rosa-tdTomato/Wt1Cre reporter mice, revealed no changes in the total number of Tomato^+^ cells, but a significant increase in Tomato^-^ cells in the heart of *Hif2/Wt1* reporter cKO mice after exposure to 2 weeks of hypoxia (Figure 6F, 6G). Furthermore, despite of the endothelial proliferation observed at 2 weeks of hypoxia (Figure 6A) in the mutant mice, FACS analyses showed no significant changes in the total number of ECs between *Hif2/Wt1* reporter cKO and the reporter control mice in hypoxia (Figure 6H, 6I, 6J). Nevertheless, there is a trend to increase on the total Tomato^-^-ECs of the *Hif2/Wt1* reporter cKO in hypoxia (Figure 6H). In addition, immunofluorescence combining endothelial (ERG, IB4) and mitosis (Ki67) markers together with endogenous Tomato signal of the reporter, revealed that the ECs proliferating in the *Hif2/Wt1* reporter cKO mice are indeed Tomato^-^ (Figure S4B). Similarly, the total PCs from *Hif2/Wt1* reporter cKO did not change significantly in hypoxia, while the control Tomato^+^-PCs were reduced. The Tomato^-^-PCs population did not change in cKO nor control mice (Figure 6K, L, M).

In sum, these results indicate that proliferation of cardiac microvascular ECs is increased after 2 weeks of hypoxia and that HIF2 signaling prevents excessive proliferation in sustained low oxygen conditions, suggesting a protective role of HIF2 against microvascular remodeling and instability in the heart during chronic hypoxia.

### Cardiopulmonary structural defects of the Hif2/Wt1 cKO are partially restored upon 1-week reoxygenation

Since the *Hif2/Wt1* cKO mice did not display alterations in basal conditions, next we wondered if the cardiac and pulmonary functional and structural defects developed after chronic hypoxia were reversible. First, we analyzed the impact of 1-week reoxygenation on 2- and 3-weeks hypoxia-induced cardiomegaly (Figure 7A), finding that both, control and *Hif2/Wt1* cKO mice, were able to restore normoxic values of heart weight to body weight ratio (Figure 7B). Moreover, reoxygenation after chronic hypoxia could also correct capillary caliber (Figure 7C) and density (Figure 7D), significantly restoring cardiac microvasculature stability. Next, we performed echocardiography analysis and determined that hypoxia-induced right (Figure 7E) and specially left (Figure 7F) ventricular hypertrophy returned to normoxia values in the *Hif2/Wt1* cKO mutant mice after 1-week reoxygenation. Furthermore, lung ultrasound revealed restoration of the pulmonary structure on *Hif2/Wt1* cKO mutants after reoxygenation (Figure 7G), with normalization of the pleural thickness and abnormal B/Z bands pattern observed after 3 weeks of chronic hypoxia (Figure 3C). The lung ultrasound analysis was further validated by classical histology showing almost complete rescue (Figure 7H) of the pulmonary congestion developed by *Hif2/Wt1* cKO mice after 3 weeks of sustained hypoxia (Figure 3D-F).

**Figure 7.**
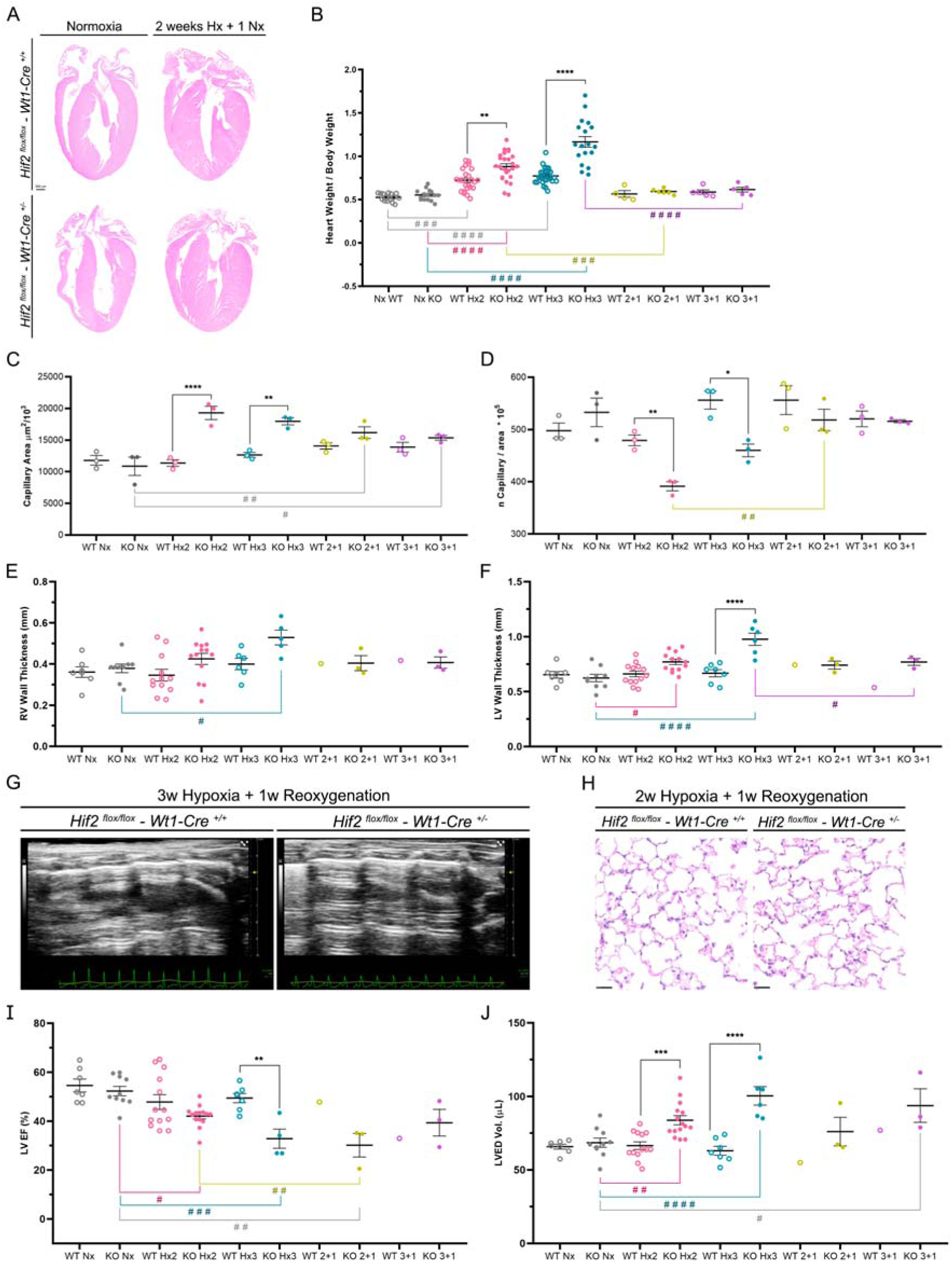
Partial rescue of chronic hypoxia-induced cardiac and pulmonary defects upon reoxygenation. **A)** Histological analysis by HE of representative heart sections of control (Hif2^flox/flox^- Wt1-Cre^+/+^, top panels) and *Hif2/Wt1* cKO (Hif2^flox/flox^-Wt1-Cre^+/-^, bottom panels) mice in normoxia (left columns) and 2 weeks of hypoxia followed by 1-week reoxygenation (2 weeks Hx+ 1Nx, right columns). Scale bar 500μm. **B)** Quantification of the heart size estimation represented as heart weight to body weight ratio in control (WT) and *Hif2/Wt1* cKO (KO) mice in normoxia (Nx, gray dots), 2 (Hx2, pink dots) or 3 weeks of Hx (Hx3, blue dots), and 2 or 3 weeks of Hx followed by 1-week reoxygenation [2+1 (yellow dots) and 3+1 (purple dots) respectively]. **C-D)** Scatter dot plot of capillary area (**C**) and density (**D**) in control (WT) and *Hif2/Wt1* cKO (KO) mice in mice in normoxia (Nx, gray dots), 2 (Hx2, pink dots) or 3 weeks of Hx (Hx3, blue dots), and 2 or 3 weeks of Hx followed by 1-week reoxygenation [2+1 (yellow dots) and 3+1 (purple dots) respectively]. **E, F)** Quantification of echocardiography analysis to asses cardiac hypertrophy by right ventricular (RV) (**E**) and left ventricular (LV) (**F**) wall thickness of control (WT) and *Hif2/Wt1* cKO (KO) mice in normoxia (Nx, gray dots), 2 (Hx2, pink dots) or 3 weeks of Hx (Hx3, blue dots), and 2 or 3 weeks of Hx followed by 1-week reoxygenation [2+1 (yellow dots) and 3+1 (purple dots) respectively]. **G)** Representative images of pulmonary ultrasound in control (Hif2^flox/flox^-Wt1-Cre^+/+^, left panel) and *Hif2/Wt1* cKO mice (Hif2^flox/flox^-Wt1-Cre^+/-^, right panel) after 3 weeks hypoxia followed by 1-week reoxygenation. **H)** Histological analysis by HE of representative lung sections in control (Hif2^flox/flox^-Wt1-Cre^+/+^, left panel) and *Hif2/Wt1* cKO mice (Hif2^flox/flox^-Wt1-Cre^+/-^, right panel) after 2 weeks hypoxia and 1-week reoxygenation. Scale bars 25μm. **I, J)** Functional and structural cardiac parameters assessed by echocardiography of control (WT) and *Hif2/Wt1* cKO (KO) mice in normoxia (Nx, gray dots), 2 (Hx2, pink dots) or 3 weeks of Hx (Hx3, blue dots), and 2 or 3 weeks of Hx followed by 1-week reoxygenation [2+1 (yellow dots) and 3+1 (purple dots) respectively]. Left ventricular ejection fraction (LVEF) (**I**) and Left ventricular end diastolic volume (LVED vol) (**J**). For all scatter plots individual values are shown and the black line represents the Mean ± SEM. Anova test. P: pvalue. * P ≤ 0,05; ** P ≤ 0,01; *** P ≤ 0,001; **** P ≤ 0,0001.

Nevertheless, despite of these structural improvements on the heart of *Hif2/Wt1* cKO mice upon 1-week reoxygenation after 2 and 3 weeks of hypoxia, functional parameters were not fully recovered, and the *Hif2/Wt1* mutants still displayed reduced left ventricular ejection fraction (Figure 7I) and elevated left ventricular end diastolic volume (Figure 7J), reflecting compromised systolic function and increased cardiac dilatation respectively.

Altogether, these results demonstrate that HIF2 signaling in the Wt1 lineage is important to maintain the correct cardiopulmonary function and structure of the microvasculature after chronic hypoxia and that the pathological changes of mice deficient for HIF2 in Wt1-contributed vascular compartments are hypoxia-dependent and reversible in contact with normal O_2_ tensions.

## Discussion

Sustained hypoxia occurring in patients with cardiorespiratory diseases and at high altitude is associated with profound vascular remodeling due to muscularization of the small arteries of the alveolar wall and proliferation of cells expressing SMA, followed by thickening of the precapillary pulmonary arteries, inflammation and fibrosis of the large proximal pulmonary arteries, leading to arterial occlusion and elevation of the RVSP (43). It has been demonstrated that this vascular remodeling and cardiac overload also occurs in rodents, allowing the use of experimental animals exposed to low oxygen as a reliable model to study PAH and cardiovascular alterations during chronic hypoxia. Endothelial HIF2 signaling has been extensively implicated in the progression of arterial remodeling leading to elevated RVSP during sustained hypoxia (18-20, 22), while deletion of HIF2 in SMCs or PCs does not prevent vascular muscularization (22, 23).

In this study, we characterized a novel mouse model, *Hif2/Wt1* cKO, to evaluate the role of HIF2 during chronic hypoxia in several vascular cell types of the cardiopulmonary system, including ECs, PCs, VSMCs and FBs (Figure 1G-M). This lineage contribution allows us to investigate the combined impact of HIF2 signaling in pulmonary and cardiac vascular cell populations beyond the endothelium by simultaneously deleting HIF2 in different cell types of these vascular compartments. Interestingly, the phenotype developed by our *Hif2/Wt1* cKO mice in response to sustained hypoxia could anticipate some effects of systemic abrogation of HIF2 with specific small molecule inhibitors that have been recently proposed for PAH treatment (33, 34). We demonstrated that HIF2 signaling in the Wt1 lineage is not necessary for proper development of the heart or lungs, neither for organ homoeostasis of adult mice (Figure S2). Furthermore, in agreement with former works, we also observed that elimination of HIF2 in pulmonary vasculature prevents arterial muscularization and elevation of right ventricular overload (Figure 2). However, the lack of HIF2 in the Wt1 lineage compromises the proper adaptation of the heart and lungs to chronic hypoxia, resulting in microvascular remodeling and dysfunction of both organs. In the lungs, we observed severe hemorrhages, inflammation and congestion that might be caused by an increased venous return associated with reduced LV ejection fraction and increased LV dilatation (Figure 5), as well as by local microvascular instability (Figure 3, 4). In the heart, despite of the absence of a hemorrhagic phenotype, mutant mice display cardiomegaly, ventricular hypertrophy without significant changes in the cardiomyocyte area, and cardiac dysfunction (Figure 4), most likely due to EC dysfunction associated with capillary dilation (Figure 5). Interestingly, most of these structural defects are restored upon reoxygenation of hypoxia-exposed *Hif2/Wt1* cKO mice, suggesting the existence of inhibitory protective roles mediated by HIF2 signaling in the cardiopulmonary vasculature only in hypoxia (Figure 7). Based on these results we proposed a working model highlighting the importance of vascular HIF2 signaling for proper capillary stability during cardiac and pulmonary adaptation to chronic hypoxia (Figure 8).

**Figure 8.**
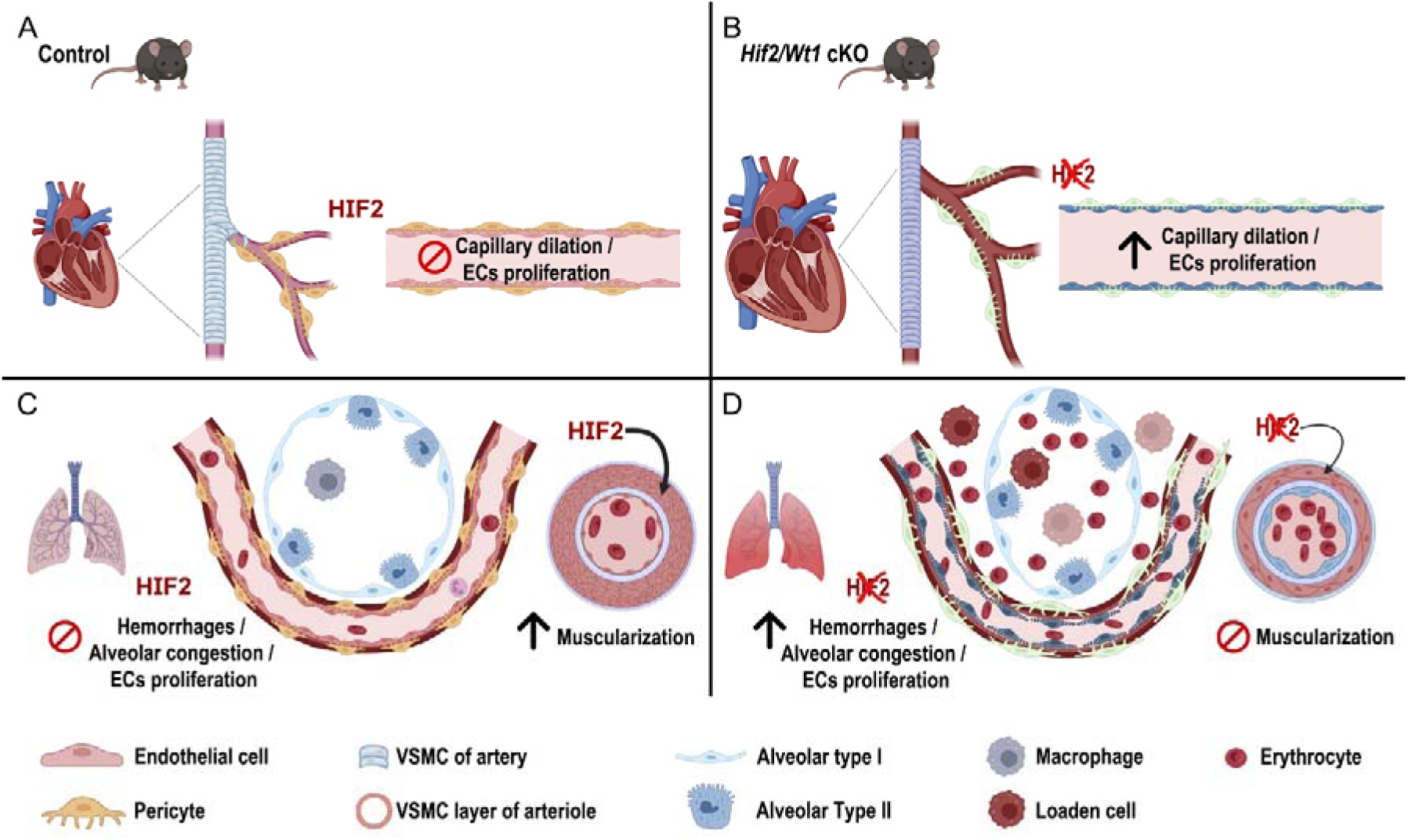
Working model. Proposed model of cardiopulmonary defects in control and *Hif2/Wt1* cKO mice after 3 weeks of chronic hypoxia. Control mice after 3 weeks of hypoxia exposure are protected against cardiomegaly, dilation of capillaries and increased proliferation of ECs (**A**), while *Hif2/Wt1* cKO mice develop cardiomegaly associated with capillary dilation and ECs proliferation (**B**). Control mice lungs display normal alveolar parenchyma as wild type HIF2 prevents microvascular proliferation, but show HIF2-mediated muscularization of the distal arterioles, and increased RVSP (**C**). In contrast, *Hif2/Wt1* cKO mice are protected against arterial muscularization, but present several structural lung defects, including erythrocytes and macrophages congestion, hemorrhages and alveolar wall thickening, probably associated to alveolar capillary remodeling and proliferation (**D**). Altogether these results suggest that HIF2 plays an inhibitory role upon cardiac and pulmonary ECs, PCs and VSMCs that prevents excessive microvascular remodeling and organ dysfunction in response to chronic hypoxia.

HIF2 global deletion protects against RVSP elevation in response to low oxygen (44) and later works demonstrated the critical role of EC-HIF2 signaling mediating the vascular remodeling that leads to elevation of RVSP during chronic hypoxia and in PAH (18-20). However, little is known about the role of HIF2 in other cell types within the vascular compartment. It has been reported that elimination of HIF2 in SMCs does not prevent RVSP elevation during chronic hypoxia (22). A recent publication by Hyunbum Kim et al. has reported novel mouse models of HIF2 overexpression and deletion in PCs using NG2-CreERT2 (23). Interestingly, while overexpression of HIF2 in PCs leads to elevation of arterial muscularization and increased RVSP after 3 weeks of hypoxia exposure - pointing to an important role of HIF2 signaling from PC mural cells in arterial vascularization-elimination of Hif2 in PCs does not prevent the elevation of RVSP during chronic hypoxia (23). These results contrast with our findings showing that the novel *Hif2/Wt1* cKO mice generated in our lab is protected against arterial remodeling and elevation of RVSP after hypoxia (Figure 2). Since Wt1 contributes to ECs, PCs, as well as SMCs and FB-like cells of the lung vasculature (Figure 1), one explanation could be that the protection of *Hif2/Wt1* cKO mice against vascular remodeling in hypoxia was only due to the ablation of HIF2 signaling in the endothelium, which would fully agree with former works (18-20). Another possibility could be that HIF2 signaling in FB-like cells of the alveolar parenchyma contributed by Wt1 also participates in the arterialization upon hypoxia. Nevertheless, Wt1 contributes to less than 4% of total pulmonary FBs (Table 2) and clarifying the precise function of HIF2 in this cell type using alternative mouse models is out of the scope of this manuscript.

Despite of the protection against arteriolar remodeling, the *Hif2/Wt1* cKO mice characterized here displayed profound damage of lung parenchyma after 3 weeks of hypoxia, with reduced alveolar space, inflammation and hemorrhages, likely due to unstable capillaries and increased remodeling of alveolar microvasculature (Figure 3). These pulmonary tissue alterations have not been reported to the best of our knowledge in the EC-specific models of HIF2 deletion. However, Hyunbum and colleagues have recently reported that overexpression of HIF2 in PCs induces vascular leakage and disruption of capillary integrity, while the PC-HIF2 KO shows no difference in permeability compared with controls (23). Taking these results into account, and considering that *Hif2/Wt1* cKO mice developed significant vascular leakage, alveolar hemorrhages and pulmonary congestion (Figure 3), together with increased EC proliferation, our results suggest that only simultaneous deletion of HIF2 in SMC/PCs and ECs lead to alveolar vascular defects and lung edema. Alternatively, we cannot rule out that HIF2 deletion in a FB-like interstitial cell type within the alveoli contributed by Wt1 could be responsible for the EC barrier dysfunction observed in our mutant.

While significant works have evaluated the relevance of HIF2 during the adaptation to sustained hypoxia in the lungs, little is known about HIF2 signaling in the heart in response to chronic low oxygen. In the setting of PAH, Smith and colleagues (45) explored the possibility that right ventricular hypertrophy after chronic hypoxia might be due not only to the increased load of the right side of the heart upon pulmonary arterial remodeling, but also to pulmonary-independent factors acting directly on the heart that could lead to activation of HIFs. Indeed, the group of Schumacker demonstrated that both, HIF1 and HIF2 signaling on cardiomyocytes, is involved in right ventricular hypertrophy during chronic hypoxia (45). In the *Hif2/Wt1* cKO mutant described here, we also demonstrated direct cardiac alterations most likely independent of the pulmonary defects occurring during chronic hypoxia. This statement is supported by the fact that *Hif2/Wt1* cKO mice developed not only right but also left ventricular hypertrophy, left ventricular systolic dysfunction based on the reduced LV ejection fraction, and left ventricular dilation (Figure 4). Moreover, considering that Wt1 contributes to capillaries of both, RV and LV (Figure 1), and that we did not observe significant differences between *Hif2/Wt1* cKO and control cardiomyocyte area in hypoxia (Figure 4), we hypothesize that the increased RV and LV wall thickness and cardiomegaly observed in the absence of HIF2 might be due to the profound microvascular remodeling and capillary dilation observed in these mutants (Figure 4). Interestingly, the recently reported Hif2/NG2 KO model does not exhibit ventricular hypertrophy, fibrosis or cardiac dysfunction in response to chronic hypoxia (23), suggesting that the vascular alterations developed by the *Hif2/Wt1* cKO mice might be due to the lack of functional HIF2 signaling in microvascular endothelium and not in cardiac PCs during low oxygen exposure. It is important to mention that in contrast with the cardiomyocyte-specific HIF2 mutant (45), the *Hif2/Wt1* cKO mice described here developed systolic dysfunction (Figure 4), while both mutants undergo ventricular dilation. These results highlight the importance of functional HIF2 signaling in the vascular compartment of the heart to ensure proper functional adaptation to sustained hypoxia. Hence, our data contribute to increase our understanding on the cardiac intrinsic response to hypoxia independently of pulmonary remodeling.

Several studies have demonstrated the role of endothelial HIF2 in the pathogenesis of PAH (18-20), and therapeutic targeting of HIF2 with small molecule inhibitors, such as belzutifan, have showed a beneficial effect in PAH *in vivo* (33, 34). Nevertheless, the consequences of prolonged systemic inhibition of HIF2 signaling in other organs essential for physiological homeostasis, like the heart, remains elusive. In this study we characterize the structural and functional cardiac and pulmonary abnormalities upon deletion of Hif2 in the Wt1 vascular/interstitial lineage during chronic hypoxia. Our results describe novel cardiac and pulmonary phenotypic characteristics of Hif2 abrogation and uncover unknown protective roles of HIF2 signaling in the microvasculature of the heart and lung, suggesting that HIF2 is essential to maintain stable microvascular networks in both organs during sustained exposure to low oxygen. These observations contribute to expand our knowledge on the role of HIF2 in the cardiovascular system and might be relevant to evaluate potential long-term effects in the setting of HIF2-specific inhibitory therapies for renal clear cell carcinoma or PAH, especially in patients with lung conditions that might suffer chronic hypoxia.

## Acknowledgements

This project has been supported by grants to SM-P from Universidad Francisco de Vitoria (UFV), LeDucq Foundation: 17CVD04, the Spanish Ministry of Science and Innovation (Ministerio de Ciencia e Innovación: MCIN): PID2020-117629RB-I00/AEI/10.13039/501100011033, Comunidad de Madrid (CM): S2022/BMD-7245 (CARDIOBOOST-CM) and Fundación Domingo Martínez: “Ayuda de Biomedicina 2023”. TA-G was supported by a predoctoral award granted by CM/EU and UFV: PEJD-2018-PRE/SAL-9529 and SM-P project PID2020-117629RB-I00/AEI/10.13039/501100011033, SM-T was funded by a predoctoral contract from Spanish Ministry of Science and Innovation and European Regional Development Fund: PRE-2021-099445, RC-M was funded by a contract from CM: PEJ-2021-AI/BMD-21926, BE was supported by SMP project 17CVD04, S.U-B was funded by SM-P project B2017/BMD-3875, J.A.N.A was supported by the University of California San Francisco (UCSF) Cardiovascular Research Institute (CVRI) department, the UCSF Dean’s office program and the Young Investigator Competitive Award from the International Society for Heart Research (ISHR), M.V-O was funded by a Juan de la Cierva Incorporación Grant (IJCI-2016-27698) E.O and S.M-P were supported by the Spanish National Research Council (CSIC).

The authors thank Jose Luis de La Pompa and Henar Cuervo for Wt1-Cre and NG2DsRed mouse lines respectively. We would like to acknowledge Virginia García, Eva Garrido, Raquel Álvarez and Mercedes de la Cueva for animal housing and handling, Antonio de Molina for histopathological analysis, Elena Prieto for assistance on FACS analysis and sorting and to CNIC Microscopy and Dynamic Imaging (ICTS-ReDib, cofounded by MCIN/AEI /10.13039/50110001103), Cellomics and Histology Core Facilities for technical assistance. We would also like to thank Mónica M-Belinchón of SEMOC (Servicio de Microscopía Óptica y Confocal del IIBM) for microscopy assistance in IIBM and Diana Muñoz for technical support.

## AUTHOR CONTRIBUTIONS

T.A-G performed most of the experiments, analyzed data, made the figures and helped to write the manuscript. M.V-O performed echocardiography and lung ultrasound analysis and discussed the manuscript. J.A. N-A contributed to FACS analysis and gating strategy and discussed the manuscript. S.R and E. O performed RVSP measurements and E.O discussed the manuscript, S.M-T, B.E, R.C-M and S.U-B supported experiments. S.M-P defined the concept; planned and supervised experiments; analyzed data; wrote the manuscript; designed the figures together with T.A-G and obtained funding to support the study.

## declaration of generative AI and AI-assisted technologies

The authors declare that no AI or AI-assisted technologies were applied to generate this manuscript.

## STAR METHODS

### Animal models care and housing

Hif2^flox/flox^ (Epas1^tm1Mcs^/J, stock #008407) (40), Rosa-tdTomato (B6.Cg-Gt(ROSA)26Sor^tm14(CAG-tdTomato)Hze^/J, stock #007914) and Ng2-DsRed (Tg(Cspg4-DsRed.T1)1Akik/J, stock #008241) (46) were obtained from Jax laboratories and maintained in homozygosity. Wt1Cre line was kindly provided by Dr. De La Pompa (32) and maintained in heterozygosity. All lines were grown on a C57BL/6 background. To generate the Hif2 floxed/Wt1Cre mouse line (*Hif2/*Wt1 cKO), homozygous Hif2^flox/flox^ females were crossed with heterozygous Wt1-Cre ^+/-^ to obtain Hif2^flox/flox^/Wt1-Cre ^+/-^ (*Hif2/*Wt1 cKO) and control Hif2^flox/flox^/Wt1-Cre ^+/+^ littermates. Hif2^flox/flox^/Wt1-Cre ^+/-^ were crossed with Rosa-tdTomato to generate the conditional HIF2/Wt1 reporter line. We did not find any difference on the phenotypic manifestations between Hif2/Wt1 cKO male and females. Hence, both sexes were indistinctly used for all the experiments. Mice were housed in SPF conditions at the CNIC Animal Facility. Welfare of animals used for experimental and other scientific purposes conformed to EU Directive 2010/63EU and Recommendation 2007/526/EC, enforced in Spanish law under Real Decreto 53/2013. Experiments with mice were allowed by the authorized Environmental Department of Comunidad de Madrid, Spain and the CNIC Animal Experimentation Ethics Committee with reference number: PROEX 267/19.

### Genotyping

All mice were genotyped by PCR using the following primers (Sigma Aldrich; USA and IDT Integrated DNA Technologies; USA) for *Hif2 floxed* alleles: Fw: 5’ GAGAGCAGCTTCTCCTGGAA 3’; Rv: 5’ TGTAGGCAAGGAAACCAAGG 3’, for Rosa-tdTomato we use four primers: Fw: 5’ AAGGGAGCTGCAGTGGAGTA 3’; Rv: 5’ CCGAAAATCTGTGGGAAGTC 3’ the wild type band and Fw: 5’ AAGGGAGCTGCAGTGGAGTA 3’; Rv: 5’ CTGTTCCTGTACGGCATGG 3’ to the mutant band. For Cre alleles, Wt1: Fw: 5’ TGACGGTGGGAGAATGTTAAT 3’; Rv: 5’ GCCGTAAATCAATCGATGAGT 3’.

### Hypoxia exposure and reoxygenation protocol

12 weeks old mice (male and female) were placed for 2 or 3 weeks inside a hypoxia chamber from Coylab (O_2_ Control Glove Box, 1 Person, Polymer, 220v). The hypoxia chamber was equipped with an oxygen control system (programed at 10% O_2_), a nitrogen gas regulator and an animal filtration system. After hypoxia exposure, mice were immediately analyzed by cardiac or lung echography and later on euthanized following the accepted protocols to proceed with organ extraction for tissue analysis. For reoxygenation experiments, the cages with animals exposed to 2- or 3-weeks hypoxia at 10% O_2_, were placed in normal ambient condition for an additional week before echography analysis and organ collection.

### Cardiopulmonary-Echography and analysis

Transthoracic echocardiography was blindly performed by an expert operator with a 30-MHz probe (Vevo 2100; VisualSonics; Canada). Mice were slightly anesthetized with 1-2% isoflurane in 100% oxygen, adjusted to maintain podal reflex (400-500 bpm, approximately). To assess the right ventricle (RV), tricuspid annular plane systolic excursion (TAPSE), acceleration time to ejection time pulmonary flow ratio (AT/ET) and RV wall thickness (RVWT) were measured. An apical four-chamber view was selected to obtain TAPSE, by M-mode. Pulmonary flow was acquired from a parasternal short axis view (SAX) at the level of the great vessels, using pulse wave Doppler. An angle SAX to optimized RV wall visualization was obtained to measure the wall thickness by M-mode. Additionally, standard SAX and parasternal long axis view (LAX) in B and M-mode were recorded and left ventricular (LV) dimensions and function were analyzed. LV end-diastolic and end-systolic (ED and ES, respectively) areas were traced for automatic calculation of the LVED and LVES volumes (LVEDV and LVESV, respectively), as well as the LV ejection fraction (LVEF). In addition, both sides of the lungs were longitudinally scanned. With these lateral views we analyzed the pleural pattern, the line profile and the predominant color indicative of edema to calculate the MoLUS (Mouse Lung UltraSound) score. To assess the MoLUS score, we assigned a value for each type of parameter (lung sliding [horizontal movement of the pleural], line profile [A, B], color profile [black or/and white], Z lines and pleural thickness, defects and effusion), according to its severity as previously described (42). The final score is the sum of all values for an individual. Images were analyzed off-line by a second blind operator.

### Right Ventricular Systolic Pressure measurement

Mice were anesthetized with medetomidine (1mg/kg) and ketamine (75mg/kg). RVSP was measured by closed-chest insertion of a Venofix A catheter (27G), coupled to a pressure transducer (Transpac IV), directly into the right ventricle. Hemodynamic data was recorded using a Biopac MP36R System and Biopac’s Acknowledge 4.1.0 software. For each mouse, at least 30 seconds of continuous and stable heartbeat cycles without noise were selected to obtain the average RVSP.

### Heart and lung extraction and processing

Mice were euthanized by CO_2_ inhalation following the approved protocol. Whole mount analysis to determine organ and body weights changes was performed at dissection. Samples were fixed for 1h (for endothelial markers) or overnight (for histological staining and non-endothelial markers) at 4°C in 4% PFA (RT15710, Electron Microscopy Sciences; PA; USA) or Formalin 10% (11699408, VWR chemicals; Avantor). After fixation, samples were embedded in 30% sucrose (16104, Sigma Aldrich; USA) and frozen in OCT medium (14020108926, Leica Biosystems) for later cryosection (8µm) preparation in a cryostat (CM1950, Leica), or were dehydrated and embedded in paraffin for sectioning at 4µm in a microtome (RM2155, Leica).

### Histological and immunohistochemical analysis

Histological analysis of heart and lung was performed using 4μm-thick paraffin sections. Sections were stained with hematoxylin & eosin (HE) for structural characterization. For arterial remodeling evaluation, lung sections were stained with an antibody against smooth muscle actin (SMA, ab150301, Abcam) following standard histological at the CNIC Histopathology Facility. HE and SMA stained slides were scanned using a Hamamatsu NanoZoomer 2.0RS device, and the image analysis and measurements were performed using NDP view (Hamamatsu; Japan) and ZEN 31 Blue Edition Lite. For immunostaining of paraffin sections, the samples were rehydrated and antigens were retrieved by incubation in citrate buffer pH 6 [10 mM sodium citrate (5949-29-1, Sigma Aldrich)] 20 minutes in a microwave. Sections were permeabilized with PBST 0,4% [PBS 1X + Triton TX100 (9036-19-5, Sigma Aldrich)] for 15 min shaking and blocked with blocking solution [PBST 0,1% with 5% goat serum (GS) (16210072, Life Technologies; NY; USA)] during 1 hour in a dark humid chamber. For OCT sections, samples were placed at room temperature for 1 and washed with distillated water for another hour. Upon complete removal of OCT, the sections were permeabilized and blocked as for paraffin sections. After blocking, sections were incubated with primary antibodies in blocking solution ON at 4°C. After washing 3 times with PBST 0,1%, sections were incubated with secondary antibodies and DAPI (Millipore, MA; USA) in PBS 1X for 1 hour at room temperature (RT) in darkness and mounted in Fluorescent Mounting Medium (S3023, Dako, Denmark or 15257659, Fisher Scientific). Images were acquired with Leica SP8 Navigator, Leica gated STED-3X-WLL SP8 or Stellaris confocal microscopes. The primary antibodies used in this study were: SMA (C6198 and A2547, Sigma Aldrich; ab150301, Abcam); ERG (ab110639, Abcam); NG2 (ZRB5320, Sigma Aldrich); pH3 (06-570, Sigma-Aldrich); Ki67 (50-5698-82, Invitrogen-ThermoFisher Scientific); Podoplanin (8.1.1, Hybridoma bank); SPC (ab211326, Abcam) and Rai3 (sc-390263, Santa Cruz). To label capillaries we used Isolectin GS-IB_4_ biotin (I21414, Invitrogen; Fisher Scientific) and to mark cell contour for cardiomyocyte identification we used WGA (W32466, Invitrogen-ThermoFisher Scientific). The secondary antibodies used in this study were: Goat anti-Rabbit AF488 (A11008, Life technologies); Goat anti-Rabbit AF594 (A11002, Invitrogen); Goat anti-Rabbit Cy3 (APC132C, EMD Millipore); Goat anti-Mouse AF488 (A11001, Invitrogen) and Streptavidin AF594 (S11227, ThermoFisher).

### Quantification of immunostaining

Immunofluorescence staining was quantified using ImageJ (Schneider, C. et al, 2012. https://doi.org/10.1038/nmeth.2089).

### Cell lineage labelling strategy for heart and lung populations by flow cytometry analysis

Mice were euthanized and subsequently the heart and all the lung lobes were perfused with HBSS (11550456, Gibco-Fisher Scientific) to clean the blood inside the tissues. On the one hand, the atria, valves and the large vessels of the heart were removed with surgical scissors, leaving only the ventricles for further analysis. On the other hand, the bronchi and trachea of the lungs were removed, leaving only the lobes. Once the tissues have been processed, cardiac ventricles and lobes of the lung were minced separately on iced-PBS. Minced heart was digested with Collagenase A (2.5 mg/ml, 10103586001, Sigma-Aldrich), liberase (0.25 mg/ml, 5401119001, Roche-Merck) and DNase I (100 u/ml, 04716728001, Roche-Merck). Minced lungs were digested only with liberase (0.25 mg/ml) and DNase I (100 u/ml). Both tissues were digested during 30 min in a water bath at 37°C with shaking every 5 minutes. Next, the digestion reaction was stopped by adding HBSS and the final single cell suspensions were obtained by mechanical dissociation and filtering in 70µm and subsequently in 40 µm Cell Strainer.

Before starting the cell lineage labelling, an erythrocyte lysis with RBC lysis buffer 1X (for 10X: 82,9 gr/L Ammonium Chloride NH_4_Cl; 10 gr/L Potassium Bicarbonate KHCO_3_, 2ml/L EDTA 0,5M, fill up to 1L of distilled water. pH7,2-7,4) was performed for 15 minutes on ice. RBC lysis reaction was stopped by adding FACS buffer (PBS 1X; 2,5% inactive FBS and EDTA 0,5M). Single cell suspensions were incubated for 45 min in rotation at 4°C with conjugated antibodies against CD45 (clone 30-F11; 103115, BioLegend); CD90.2 (clone 53-2.1; 140319, BioLegend); CD31 (clone 390; 102427, BioLegend); Podoplanin-Gp38 (clone 8.1.1; 127423, BioLegend) and CD39 (clone Duha59; 143805, BioLegend) to define gates for non-myocyte populations in the heart and lung as previously described by others (37, 47-49). Sytox Green was used as viability marker. Samples were acquired in LSRFortessa (BD FORTESSA SORP) and FACSAria Fusion Cell Sorter (BD FACSAria™ Fusion CellSorter (BSL-2) equipped with DIVA software. FlowJo was used to analyze the data.

### Quantification of non-myocyte cell percentage or total number per gram of tissue

To calculate the percentage of non-myocyte cells in the heart and lung contributed by the Wt1 lineage, we used Wt1-Cre/Rosa-tdTomato and followed the digestion and labeling strategy described above. To determine the contribution to each cardiac lineage, we calculated the number of each cell population within the CD45^-^/Wt1-Tomato^+^ gate relative to the total Tomato^+^ cells following the formula:

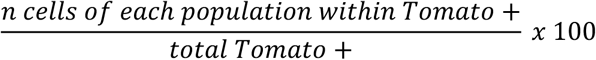

To determine the absolute number of each cell lineage contributed by Wt1 in different mice, we used beads to normalize the value per gram of tissue. In particular, we used Truecount beads (663028, BD-Biosciences) that were prepared at a 10000 beads/ml of FACS buffer with sytox green, viability marker. 500 µl of this Beads Buffer were added to the cell suspension (already labeled) described above. Approximately 1,000 beads were acquired per sample. The quantification of the number of cells per gram of tissue was calculated with this formula:

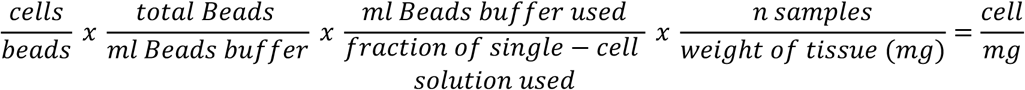

In this equation, cells/beads are the exact number of cells/beads after FlowJo analysis. Total beads/ml Beads buffer are 10000 beads/1 ml Beads buffer. 0,5 ml of Beads buffer was used per 0,1 ml of single-cell suspension of digestion tissue and per tube there are only 1 sample.

## References

1. Simon MC, and Keith B. The role of oxygen availability in embryonic development and stem cell function. Nat Rev Mol Cell Biol. 2008;9(4):285–96.

2. Kaelin WG, Jr., and Ratcliffe PJ. Oxygen sensing by metazoans: the central role of the HIF hydroxylase pathway. Mol Cell. 2008;30(4):393-402.

3. Majmundar AJ, Wong WJ, and Simon MC. Hypoxia-inducible factors and the response to hypoxic stress. Mol Cell. 2010;40(2):294-309.

4. Dengler VL, Galbraith M, and Espinosa JM. Transcriptional regulation by hypoxia inducible factors. Crit Rev Biochem Mol Biol. 2014;49(1):1-15.

5. Koh MY, and Powis G. Passing the baton: the HIF switch. Trends Biochem Sci. 2012;37(9):364–72.

6. Hu CJ, Wang LY, Chodosh LA, Keith B, and Simon MC. Differential roles of hypoxia-inducible factor 1alpha (HIF-1alpha) and HIF-2alpha in hypoxic gene regulation. Mol Cell Biol. 2003;23(24):9361-74.

7. Mole DR, Blancher C, Copley RR, Pollard PJ, Gleadle JM, Ragoussis J, et al. Genome-wide association of hypoxia-inducible factor (HIF)-1alpha and HIF-2alpha DNA binding with expression profiling of hypoxia-inducible transcripts. J Biol Chem. 2009;284(25):16767-75.

8. Knutson AK, Williams AL, Boisvert WA, and Shohet RV. HIF in the heart: development, metabolism, ischemia, and atherosclerosis. J Clin Invest. 2021;131(17).

9. Krishnan J, Ahuja P, Bodenmann S, Knapik D, Perriard E, Krek W, et al. Essential role of developmentally activated hypoxia-inducible factor 1alpha for cardiac morphogenesis and function. Circ Res. 2008;103(10):1139-46.

10. Guimaraes-Camboa N, Stowe J, Aneas I, Sakabe N, Cattaneo P, Henderson L, et al. HIF1alpha Represses Cell Stress Pathways to Allow Proliferation of Hypoxic Fetal Cardiomyocytes. Developmental cell. 2015;33(5):507-21.

11. Menendez-Montes I, Escobar B, Gomez MJ, Albendea-Gomez T, Palacios B, Bonzon-Kulichenko E, et al. Activation of amino acid metabolic program in cardiac HIF1-alpha-deficient mice. iScience. 2021;24(2):102124.

12. Menendez-Montes I, Escobar B, Palacios B, Gomez MJ, Izquierdo-Garcia JL, Flores L, et al. Myocardial VHL-HIF Signaling Controls an Embryonic Metabolic Switch Essential for Cardiac Maturation. Dev Cell. 2016;39(6):724-39.

13. Tian H, Hammer RE, Matsumoto AM, Russell DW, and McKnight SL. The hypoxia-responsive transcription factor EPAS1 is essential for catecholamine homeostasis and protection against heart failure during embryonic development. Genes Dev. 1998;12(21):3320-4.

14. Scortegagna M, Ding K, Oktay Y, Gaur A, Thurmond F, Yan LJ, et al. Multiple organ pathology, metabolic abnormalities and impaired homeostasis of reactive oxygen species in Epas1-/- mice. Nat Genet. 2003;35(4):331-40.

15. Peng J, Zhang L, Drysdale L, and Fong GH. The transcription factor EPAS-1/hypoxia-inducible factor 2alpha plays an important role in vascular remodeling. Proc Natl Acad Sci U S A. 2000;97(15):8386-91.

16. Shimoda LA, and Semenza GL. HIF and the lung: role of hypoxia-inducible factors in pulmonary development and disease. Am J Respir Crit Care Med. 2011;183(2):152-6.

17. Sheikh AQ, Saddouk FZ, Ntokou A, Mazurek R, and Greif DM. Cell Autonomous and Non-cell Autonomous Regulation of SMC Progenitors in Pulmonary Hypertension. Cell Rep. 2018;23(4):1152-65.

18. Cowburn AS, Crosby A, Macias D, Branco C, Colaco RD, Southwood M, et al. HIF2alpha-arginase axis is essential for the development of pulmonary hypertension. Proc Natl Acad Sci U S A. 2016;113(31):8801-6.

19. Kapitsinou PP, Rajendran G, Astleford L, Michael M, Schonfeld MP, Fields T, et al. The Endothelial Prolyl-4-Hydroxylase Domain 2/Hypoxia-Inducible Factor 2 Axis Regulates Pulmonary Artery Pressure in Mice. Mol Cell Biol. 2016;36(10):1584-94.

20. Dai Z, Li M, Wharton J, Zhu MM, and Zhao YY. Prolyl-4 Hydroxylase 2 (PHD2) Deficiency in Endothelial Cells and Hematopoietic Cells Induces Obliterative Vascular Remodeling and Severe Pulmonary Arterial Hypertension in Mice and Humans Through Hypoxia-Inducible Factor-2alpha. Circulation. 2016;133(24):2447-58.

21. Kurakula K, Smolders V, Tura-Ceide O, Jukema JW, Quax PHA, and Goumans MJ. Endothelial Dysfunction in Pulmonary Hypertension: Cause or Consequence? Biomedicines. 2021;9(1).

22. Tang H, Babicheva A, McDermott KM, Gu Y, Ayon RJ, Song S, et al. Endothelial HIF-2alpha contributes to severe pulmonary hypertension due to endothelial-to-mesenchymal transition. Am J Physiol Lung Cell Mol Physiol. 2018;314(2):L256-L75.

23. Kim H, Liu Y, Kim J, Kim Y, Klouda T, Fisch S, et al. Pericytes contribute to pulmonary vascular remodeling via HIF2alpha signaling. EMBO Rep. 2024;25(2):616-45.

24. Wilm B, and Munoz-Chapuli R. The Role of WT1 in Embryonic Development and Normal Organ Homeostasis. Methods Mol Biol. 2016;1467:23–39.

25. Munoz-Chapuli R, Macias D, Gonzalez-Iriarte M, Carmona R, Atencia G, and Perez-Pomares JM. [The epicardium and epicardial-derived cells: multiple functions in cardiac development]. Rev Esp Cardiol. 2002;55(10):1070-82.

26. Martinez-Estrada OM, Lettice LA, Essafi A, Guadix JA, Slight J, Velecela V, et al. Wt1 is required for cardiovascular progenitor cell formation through transcriptional control of Snail and E-cadherin. Nat Genet. 2010;42(1):89-93.

27. Zhou B, Ma Q, Rajagopal S, Wu SM, Domian I, Rivera-Feliciano J, et al. Epicardial progenitors contribute to the cardiomyocyte lineage in the developing heart. Nature. 2008;454(7200):109–13.

28. Wagner N, Ninkov M, Vukolic A, Cubukcuoglu Deniz G, Rassoulzadegan M, Michiels JF, et al. Implications of the Wilms’ Tumor Suppressor Wt1 in Cardiomyocyte Differentiation. Int J Mol Sci. 2021;22(9).

29. Diaz Del Moral S, Barrena S, Hernandez-Torres F, Aranega A, Villaescusa JM, Gomez Doblas JJ, et al. Deletion of the Wilms’ Tumor Suppressor Gene in the Cardiac Troponin-T Lineage Reveals Novel Functions of WT1 in Heart Development. Front Cell Dev Biol. 2021;9:683861.

30. Duim SN, Goumans MJ, and Kruithof BPT. In: van den Heuvel-Eibrink MM ed. Wilms Tumor. Brisbane (AU); 2016.

31. Cano E, Carmona R, and Munoz-Chapuli R. Wt1-expressing progenitors contribute to multiple tissues in the developing lung. Am J Physiol Lung Cell Mol Physiol. 2013;305(4):L322-32.

32. del Monte G, Casanova JC, Guadix JA, MacGrogan D, Burch JB, Perez-Pomares JM, et al. Differential Notch signaling in the epicardium is required for cardiac inflow development and coronary vessel morphogenesis. Circ Res. 2011;108(7):824-36.

33. Macias D, Moore S, Crosby A, Southwood M, Du X, Tan H, et al. Targeting HIF2alpha-ARNT hetero-dimerisation as a novel therapeutic strategy for pulmonary arterial hypertension. Eur Respir J. 2021;57(3).

34. Hu CJ, Poth JM, Zhang H, Flockton A, Laux A, Kumar S, et al. Suppression of HIF2 signalling attenuates the initiation of hypoxia-induced pulmonary hypertension. Eur Respir J. 2019;54(6).

35. Zhou B, and Pu WT. Genetic Cre-loxP assessment of epicardial cell fate using Wt1-driven Cre alleles. Circ Res. 2012;111(11):e276-80.

36. Horie M, Castaldi A, Sunohara M, Wang H, Ji Y, Liu Y, et al. Integrated Single-Cell RNA-Sequencing Analysis of Aquaporin 5-Expressing Mouse Lung Epithelial Cells Identifies GPRC5A as a Novel Validated Type I Cell Surface Marker. Cells. 2020;9(11).

37. Travaglini KJ, Nabhan AN, Penland L, Sinha R, Gillich A, Sit RV, et al. A molecular cell atlas of the human lung from single-cell RNA sequencing. Nature. 2020;587(7835):619–25.

38. Lin C, Song H, Huang C, Yao E, Gacayan R, Xu SM, et al. Alveolar type II cells possess the capability of initiating lung tumor development. PLoS One. 2012;7(12):e53817.

39. Sorokin SP, and Hoyt RF, Jr. Macrophage development: I. Rationale for using Griffonia simplicifolia isolectin B4 as a marker for the line. Anat Rec. 1992;232(4):520-6.

40. Gruber M, Hu CJ, Johnson RS, Brown EJ, Keith B, and Simon MC. Acute postnatal ablation of Hif-2alpha results in anemia. Proc Natl Acad Sci U S A. 2007;104(7):2301-6.

41. Sheikh AQ, Misra A, Rosas IO, Adams RH, and Greif DM. Smooth muscle cell progenitors are primed to muscularize in pulmonary hypertension. Sci Transl Med. 2015;7(308):308ra159.

42. Villalba-Orero M, Lopez-Olaneta MM, Gonzalez-Lopez E, Padron-Barthe L, Gomez-Salinero JM, Garcia-Prieto J, et al. Lung ultrasound as a translational approach for non-invasive assessment of heart failure with reduced or preserved ejection fraction in mice. Cardiovasc Res. 2017;113(10):1113-23.

43. Stenmark KR, Meyrick B, Galie N, Mooi WJ, and McMurtry IF. Animal models of pulmonary arterial hypertension: the hope for etiological discovery and pharmacological cure. Am J Physiol Lung Cell Mol Physiol. 2009;297(6):L1013-32.

44. Brusselmans K, Compernolle V, Tjwa M, Wiesener MS, Maxwell PH, Collen D, et al. Heterozygous deficiency of hypoxia-inducible factor-2alpha protects mice against pulmonary hypertension and right ventricular dysfunction during prolonged hypoxia. J Clin Invest. 2003;111(10):1519-27.

45. Smith KA, Waypa GB, Dudley VJ, Budinger GRS, Abdala-Valencia H, Bartom E, et al. Role of Hypoxia-Inducible Factors in Regulating Right Ventricular Function and Remodeling during Chronic Hypoxia-induced Pulmonary Hypertension. Am J Respir Cell Mol Biol. 2020;63(5):652-64.

46. Zhu X, Bergles DE, and Nishiyama A. NG2 cells generate both oligodendrocytes and gray matter astrocytes. Development. 2008;135(1):145-57.

47. Skelly DA, Squiers GT, McLellan MA, Bolisetty MT, Robson P, Rosenthal NA, et al. Single-Cell Transcriptional Profiling Reveals Cellular Diversity and Intercommunication in the Mouse Heart. Cell Rep. 2018;22(3):600-10.

48. Stellato M, Czepiel M, Distler O, Blyszczuk P, and Kania G. Identification and Isolation of Cardiac Fibroblasts From the Adult Mouse Heart Using Two-Color Flow Cytometry. Front Cardiovasc Med. 2019;6:105.

49. Ugorski M, Dziegiel P, and Suchanski J. Podoplanin - a small glycoprotein with many faces. Am J Cancer Res. 2016;6(2):370-86.

