## Supplemental figures and figure legends for "Vascular HIF2 signaling prevents cardiomegaly, alveolar congestion and capillary remodeling during chronic hypoxia"

### Supplemental material

#### *Figure S1*

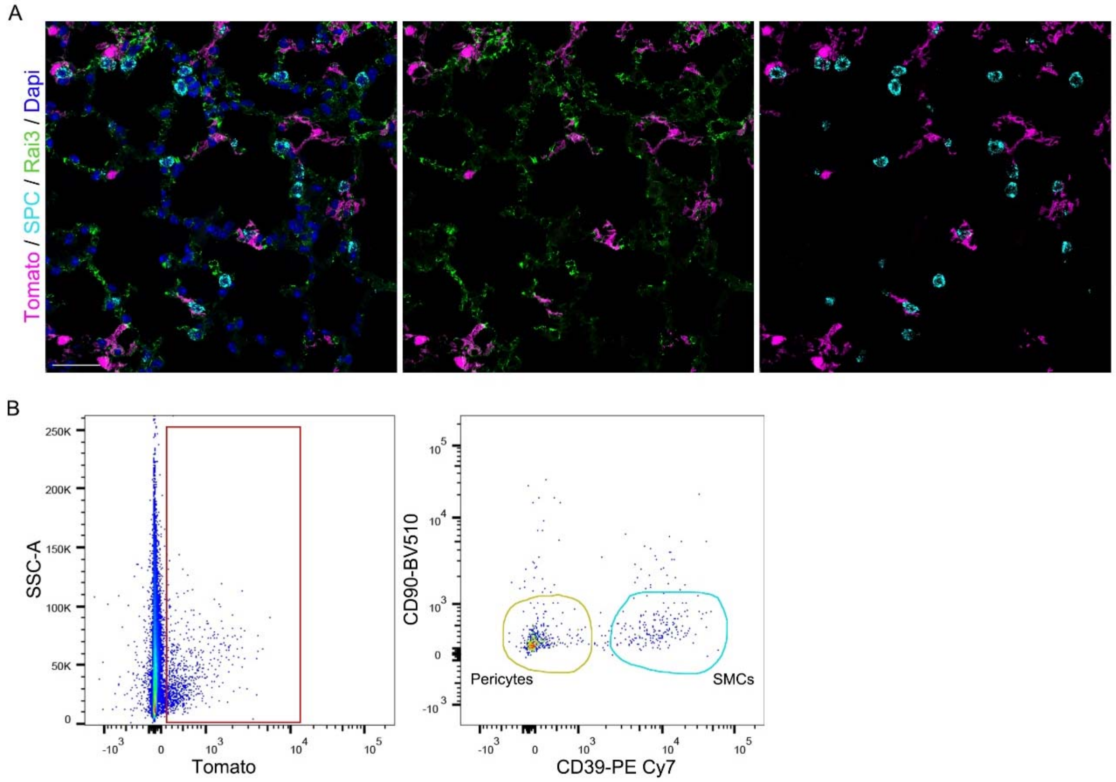


**Figure S1. Wt1 lineage tracing in adult lung tissue. A)** Immunofluorescence of representative lung sections of Rosa-tdTomato/Wt1Cre reporter mice with cell lineage markers for alveolar type I (Rai3, green) and alveolar type II [Surfactant Protein-C (SPC), cyan] cells and Tomato (Wt1 lineage, magenta). Nucleus are co-stained with Dapi (blue). Scale bar 40μm. **B)** FACs analysis plot of the NG2-DsRed reporter mice. Left panel show the side scatter area (SSC-A) versus Tomato signal and the red square represents the Tomato$^+$ cells. The right panel shows the distribution of PCs (yellow gate) and SMCs (blue gate) from Tomato$^+$ cells (left) when stained with CD90 and CD39. The yellow gate for PCs applied in the Wt1/Tomato lineage analysis overlaps with the one of NG2-DsRed, confirming the adequacy of this strategy for PC identification.

*Figure S2*

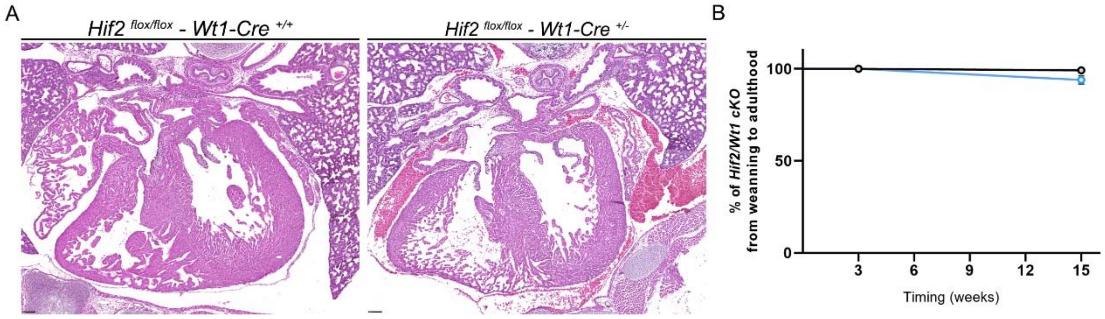


**Figure S2. Embryonic phenotype and survival curve of *Hif2/Wt1* cKO mice. A)** Histological analysis by HE of embryonic E18.5 heart sections from control (Hif2 $^{flox/flox}$-Wt1-Cre $^{+/+}$, left panel) and *Hif2/Wt1* cKO (Hif2 $^{flox/flox}$-Wt1-Cre $^{+/-}$, right panel) embryos. Scale bars 100μm.**B)** Survival curve showing the percentage of control (black line) and *Hif2/Wt1* cKO (blue line) mice recovered from weaning (3 weeks) to adulthood (up to 15 weeks).

*Figure S3*

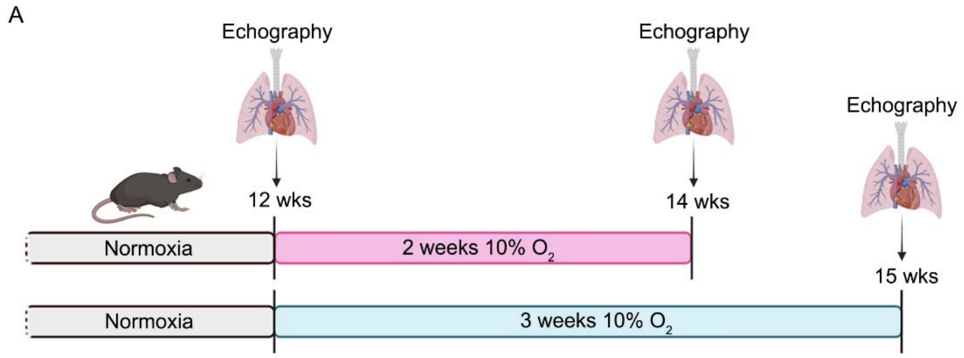


**Figure S3. Experimental workflow.** Schematic of the experimental design. 12 weeks old mice were place into a hypoxia chamber at 10% oxygen for 2 (pink line) or 3 (blue line) weeks. After hypoxia exposure, mice were analyzed by cardiac and lung echography and then, euthanized following the accepted protocols to proceed with organ extraction for tissue analysis.

*Figure S4*

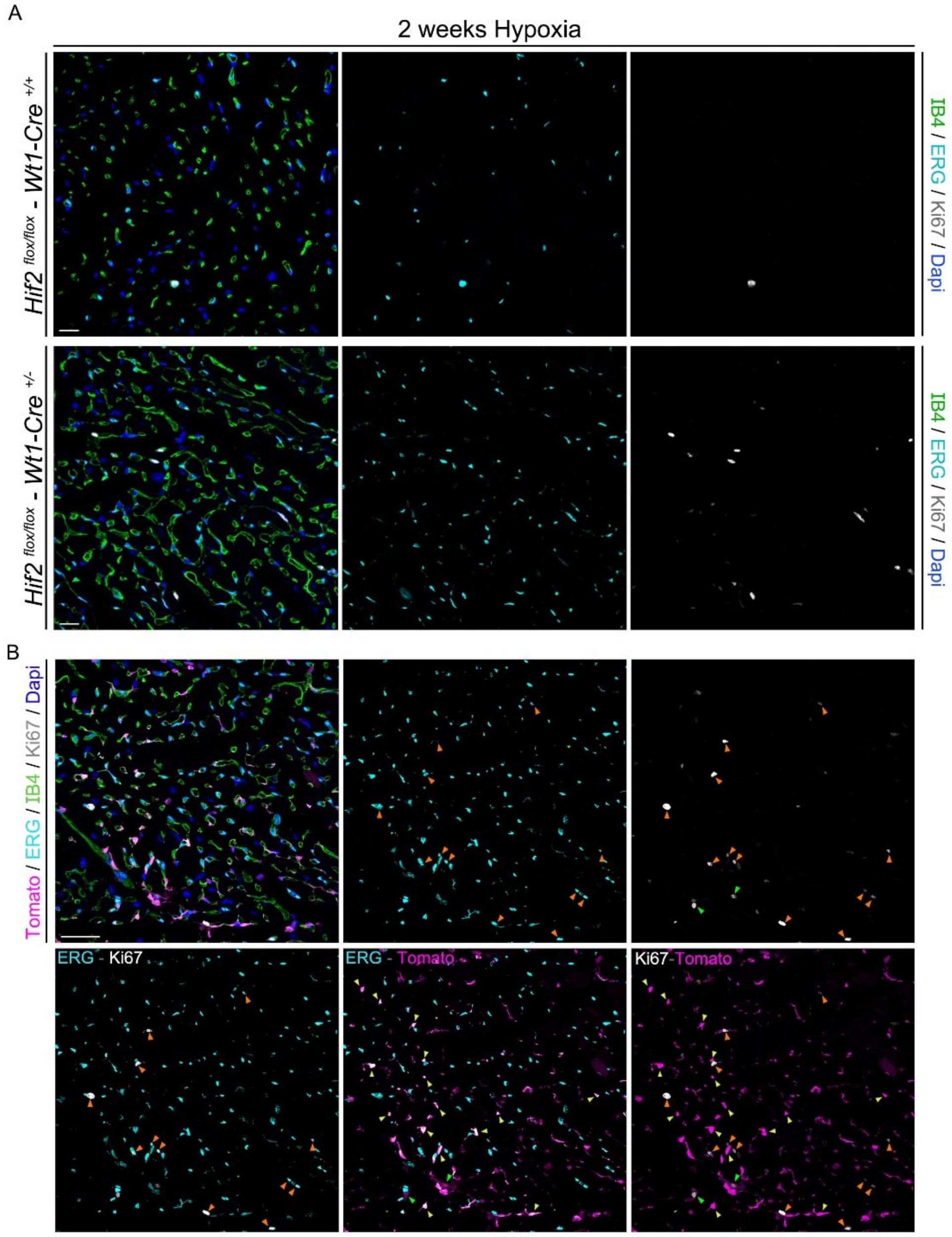


**Figure S4. Proliferation of Wt1-derived cardiac ECs during chronic hypoxia. A)** Immunofluorescence with IB4 (outline of ECs, green), ERG (nucleus of ECs, cyan), Ki67 (mitosis, white) and Dapi (nucleus, blue) of representative cardiac sections of control (Hif2$^{flox/flox}$-Wt1-Cre$^{+/+}$, top panels) and *Hif2/Wt1* cKO (Hif2$^{flox/flox}$-Wt1-Cre$^{+/-}$, bottom panels) mice after 2 weeks of hypoxia. Scale bars 20μm. **B)** Representative ERG (ECs, cyan), Ki67 (mitosis, white) Tomato (Wt1 lineage, magenta), IB4 (outline of ECs, green) and nucleus (Dapi, blue) staining of cardiac sections from *Hif2/Wt1* cKO mice. Merge panels show no localization between Tomato and Ki67. Orange arrowheads indicate ERG$^{+}$/Ki67$^{+}$/Tomato$^{-}$ ECs. Yellow arrowheads indicate ERG$^{+}$/Ki67$^{-}$/Tomato$^{+}$ ECs. Green arrowheads indicate a Wt1-lineage non-endothelial proliferating cell, probably a PC. Scale bar 40μm.
